# A multi-omic dissection of super-enhancer driven oncogenic gene expression programs in ovarian cancer

**DOI:** 10.1101/2022.04.08.487699

**Authors:** Michael R. Kelly, Kamila Wisniewska, Matthew J. Regner, Michael W. Lewis, Andrea A. Perreault, Eric S. Davis, Douglas H. Phanstiel, Joel S. Parker, Hector L. Franco

## Abstract

The human genome contains regulatory elements, such as enhancers, that are often rewired by cancer cells for the activation of genes that promote tumorigenesis and resistance to therapy. This is especially true for cancers that have little or no known driver mutations within protein coding genes, such as ovarian cancer. Herein, we have utilized an integrated set of genomic and epigenomic datasets to identify clinically relevant super-enhancers that are preferentially amplified in ovarian cancer patients. We have systematically probed the top 86 super-enhancers, using CRISPR-interference and CRISPR-deletion assays coupled to RNA-sequencing, to nominate two salient super-enhancers that drive proliferation and migration of cancer cells. Utilizing Hi-C, we constructed chromatin interaction maps that enabled the annotation of direct target genes for these super-enhancers and later confirmed their activity specifically within the cancer cell compartment of human tumors using single-cell genomics data. Together, our multi-omic approach has examined a number of fundamental questions about how regulatory information encoded into super-enhancers drives gene expression networks that underlie the biology of ovarian cancer.

## INTRODUCTION

Ovarian cancer is one of the deadliest cancers among women worldwide and is the leading cause of gynecologic-related cancer deaths in the U.S.^4^. High grade serous ovarian cancer (HGSOC) is the most common subtype (approximately 80% of all ovarian cancer) and is characterized by a high number of copy number alterations and few driver mutations, which is thought to account for the clinical aggressiveness of this disease as well as the eventual development of chemoresistance^5, 6^. The most commonly seen mutation in HGSOC is p53 (>90% of cases) followed by low prevalence but statistically significant recurrent somatic mutations in NF-1, BRCA1/2, and CDK2, which often lead to genomic instability^7–9^. Due to this genomic instability, ovarian cancer has a high rate of copy number abnormalities and recent studies have shown that these alterations can be used to stratify HGSOC^5^. However, the paucity of known driver mutations for ovarian cancer has made it difficult to develop effective targeted therapies. Consequently, the standard of care remains cytoreductive surgery followed by carboplatin/taxane chemotherapy, with approximately 75% patients experiencing a recurrence^110^. Thus, additional analysis of the non-coding regions of the genome, that extends beyond gene profiling, is desperately needed.

Mounting evidence suggests that regulatory elements, such as transcriptional enhancers, can be rewired or hijacked by cancer cells for the activation of genes that promote tumor formation, metastasis, and resistance to therapy^1–3^. This is especially true for cancers that have little or no known driver mutations within protein coding genes, such as ovarian cancer^11^. Enhancers are non-coding DNA elements that contain information for the binding of transcription factors and interact spatially with their target genes to orchestrate spatiotemporal patterns of gene expression^12, 13^. It is estimated that there are hundreds of thousands of enhancers found throughout our genome and these can act independent of orientation and linear distance from their target genes, forming high order chromatin loops with their target genes. Of note, the activity of enhancers is often restricted to a particular cell type or specific physiological or pathological conditions, enabling their genomic function to determine precisely when, where, and at what level each of our genes is expressed^14–16^. Large clusters of neighboring enhancers that have unusually high occupancy of interacting factors are typically called super-enhancers (SEs)^17^. These super-enhancers are known to regulate key cell identity genes, and in cancer are known to drive oncogene expression^18^.

The high transcriptional output of cancer cells is thought to be sustained by the activity of super-enhancers, suggesting cancer cells can become addicted to super-enhancer driven regulatory networks^19^. Furthermore, recent studies in ovarian cancer have demonstrated the capacity of super-enhancers and their associated networks of transcription factors to directly influence chemoresistance^20, 21^. The molecular characteristics and high activity of super-enhancers make them exquisitely sensitive to epigenetic drugs, more so than typical enhancers^22^. Thus, there is a growing belief that exploiting transcriptional dependence by targeting oncogenic super-enhancers may be a valid therapeutic avenue^22^. For example, the Bromodomain Containing Protein 4 (BRD4) is a druggable transcription factor that recognizes acetylated histone proteins and is found in large quantities at super-enhancers^23, 24^. Small molecule inhibition of BRD4 (such as JQ1 and BET inhibitors) has been shown to reduce cell proliferation and survival *in vivo* as well as increase therapeutic sensitivity of several cancer types, leading to the development of several clinical trials^21, 24, 25^. However, despite their effectiveness in inhibiting oncogenic processes in ovarian cancer cells, anti-BRD4 agonists remain a poor therapeutic option due to their overall toxicity and delivery constraints^26^. Nevertheless, the study of BRD4 associated super-enhancers in ovarian cancer may lead to the identification of biomarkers, downstream druggable targets, and a better understanding of the regulatory processes that drive this disease.

To this end, the studies described herein have examined several fundamental questions about how regulatory information is encoded into super-enhancers, how they are preferentially amplified in ovarian cancer cells, and how they drive gene expression networks that underlie the biology of ovarian cancer cells. We used an integrated genomic and computational framework to (1) identify BRD4-enriched and copy number amplified super-enhancers in ovarian cancer patients, (2) systematically probe the functions of the top 86 ovarian cancer specific super-enhancers using CRISPR interference assays (CRISPRi) (dCas9-KRAB) coupled to RNA-seq, (3) validate their roles in driving the proliferation and migration of cancer cells via CRISPR-knockouts, (4) annotate direct target genes using chromatin looping information via Hi-C, and (5) confirm their activity specifically within the cancer cell compartment of human tumors using single cell genomics data.

## RESULTS

### Identification of BRD4-Enriched Super-Enhancers in Ovarian Cancer

Super-enhancers are one of the most salient regulatory elements in the genome and are known to be repurposed by cancer cells to drive the expression of oncogenes^17, 27^. Due to the unusually high levels of interacting transcription factors and the prominence of their target genes, super-enhancers contain untapped potential that can lead to a new set of markers with diagnostic and prognostic potential, or even serve as tractable targets for therapeutic intervention^22, 28^. To identify enhancers likely to be associated with oncogenic gene expression programs, we leveraged both ovarian cancer cell line epigenetic data and patient tumor RNA-seq and Copy Number data from The Cancer Genome Atlas (TCGA) (Figure 1A).

**Figure 1.**
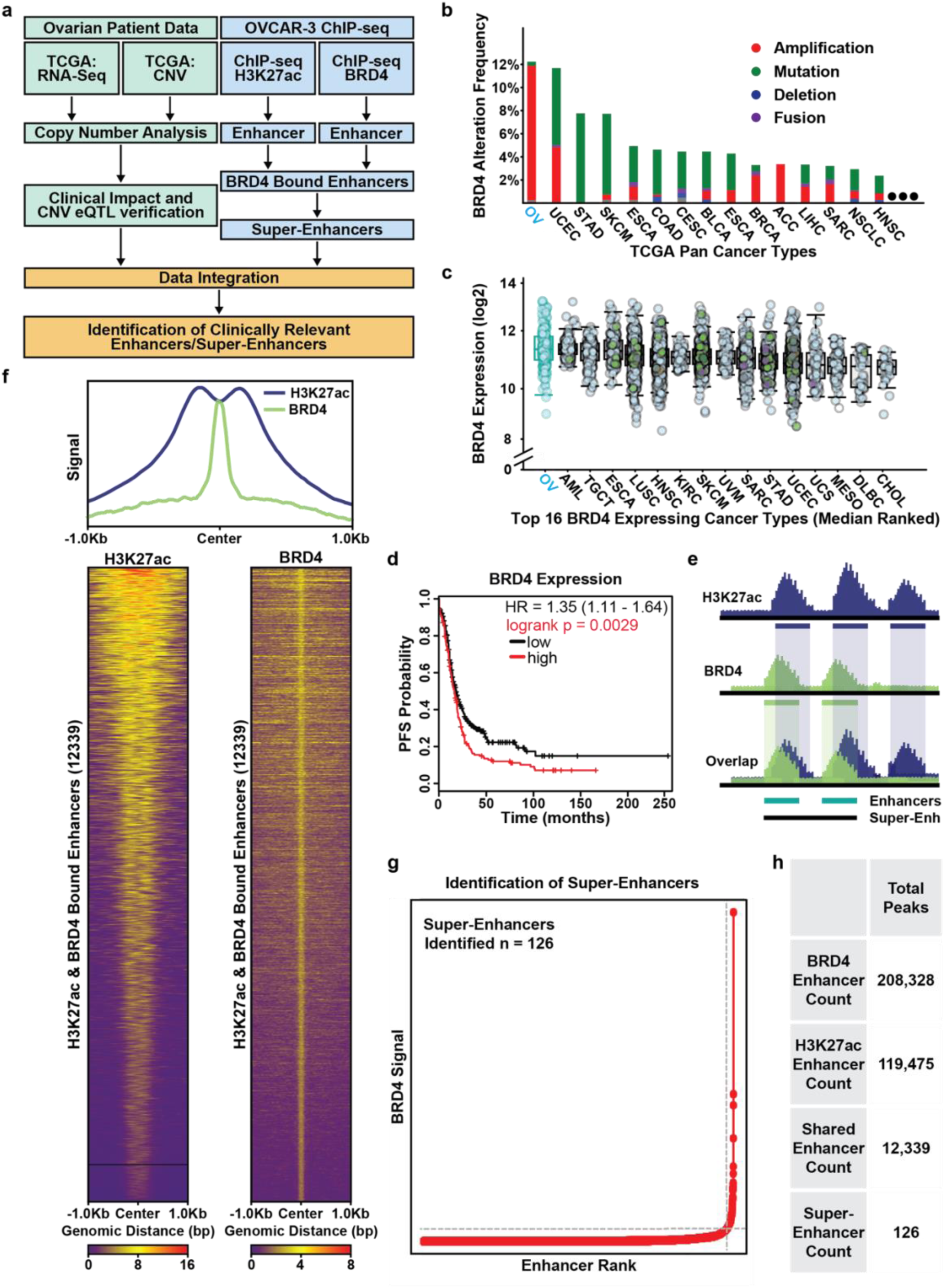
Identification of BRD4-Enriched Super-Enhancers in Ovarian Cancer. **a.** Flowchart of the analysis strategy used to identify clinically relevant BRD4-enriched SEs in ovarian cancer. **b.** Bar chart depicting the alteration frequency of the BRD4 locus across the TGCA Pan Cancer patient cohort (ovarian cancer = OV). **c.** Box plots showing normalized BRD4 expression across the top 16 highest expressing cancer types in the TCGA Pan Cancer patient cohort (ovarian cancer = OV). **d.** Kaplan-Meier plots showing the relationship between BRD4 expression and progression free survival in ovarian cancer patients with high grade serous (n = 1232) or endometrioid histology (n = 62). Patients are split by median expression of BRD4 and the red line represents patients in the high expression cohort and the black the line low expression cohort. **e.** Cartoon depicting the analysis strategy for integrating H3K27ac and BRD4 ChIP-seq data and selecting overlapping peaks to call super-enhancers. BRD4 is shown in green and H3K27ac in blue. **f.** *Top:* Meta-ChIP plot of the signal across shared peaks showing overlap of H3K27ac and BRD4 signal. *Bottom:* Heatmap of ChIP signal across all 12,339 called shared peaks. The samples are scaled relative to the background for that signal group independent of the other signal (BRD4 to BRD4 background; H3K27ac to H3K27ac background). **g.** BRD4 signal versus enhancer rank plot showing the identification of 126 super-enhancers as defined by the ROSE software. **h.** Tabulation of the total number of enhancers/peaks identified.

First, we used existing ChIP-seq data in the well vetted high-grade serous ovarian cancer cell line OVCAR3 to identify active enhancers by searching for co-localization of the histone modification histone H3 lysine 27 acetylation (H3K27ac) and BRD4 (Figure 1F)^29–31^. BRD4 enrichment was considered a critical component for the detection of potentially oncogenic enhancers due to key observations previously shown in ovarian cancer patients^24^. Namely, across the entirety of the TCGA Pan Cancer dataset, ovarian cancer patients have the highest rate of genetic alterations at the BRD4 locus, with ∼11% of patients having an amplification of this region (Figure 1B)^1, 8, 9^. Moreover, ovarian cancer has the highest overall expression of BRD4 across all TCGA cancer types and patients with increased expression of BRD4 experienced significantly reduced survival times as determined through Kaplan-Meier analysis (Figure 1C and D)^9, 32, 33^. Therefore, we defined active enhancers as intergenic regions that contained at least a 1-base pair overlap between statistically significant BRD4 peaks and H3K27ac peaks called by the MACS2 peak calling algorithm (Figure 1E)^34^. To focus on distal enhancer elements, any peaks that overlapped with annotated genes or promoter regions were removed. This pipeline identified 12,339 BRD4-enriched active enhancer elements in ovarian cancer cells. To determine if these enhancers are lineage specific or extensible to other cancer types, we investigated the overlap with existing enhancer annotations across normal tissues (defined by the ENCODE consortium) and across existing annotations in other cancer types (Supplemental Figure 1B and C)^17, 35^. We found that 44.1% of the 12,339 BRD4-enriched enhancers had at least 1-base pair overlap with active enhancers in normal tissues and this number increases to 73.6% when comparing to active enhancers across several cancer types (Supplemental Figure 1B). The aforementioned importance of BRD4 and the high degree of overlap between these enhancers with cancer-specific enhancers gave us confidence for using these data for calling super-enhancers.

From our pool of 12,339 constituent enhancers, we identified 126 super-enhancer regions using the Rank Ordering of Super-Enhancers (ROSE) algorithm (Figure 1G and H, Supplemental Data 1, 2, 3, and 4)^36, 37^. To determine if these BRD4-enriched super-enhancers are relevant to ovarian cancer patients, we leveraged single-cell assay for transposase-accessible chromatin sequencing (ATAC-seq) data generated from HGSOC patients to measure the activity of the super-enhancers within these tumors^38^. We detected the activity (defined by chromatin accessibility) of 121 out of 126 (96%) super-enhancers in the cancer cell fraction of HGSOC patients (Supplemental Figure 1A). Taken together, these data suggest that the super-enhancers identified using our pipeline are not cell line specific and may be relevant to both ovarian cancer and other cancer types. To further investigate the clinical utility of these SEs, we next looked for evidence in patient tumors using both TCGA RNA-seq and Copy Number Variation data.

### Copy Number Variation and Expression Quantitative Trait Loci (CNVeQTL) Analysis Nominate Putative Oncogenic Super-Enhancers

Given that copy number variation has been previously identified as an important hallmark of ovarian cancer, we sought to investigate whether these BRD4-enriched super-enhancers were preferentially amplified in ovarian cancer patients^5^. To this end, we performed a computational experiment making use of publicly available copy number variation data across nearly 600 ovarian cancer patients^11^ to compare the copy number amplification values overlapping our SE regions to the amplification across the ovarian cancer genome as a whole, by both random-draw (pseudo-bootstrap) and direct comparison analyses (Figure 2A). Copy number variation (CNV) values across ∼600 ovarian cancer patients were quantified by dividing the genome into uniform 15kb sliding windows and assigning CNV segment values within each window (Figure 2B). We then compared the amplification of the SE overlapping windows against an equivalent number of randomly drawn windows across the ovarian cancer genome (inclusive of our SE regions). The random drawing of windows was iterated 10,000 times and, in each comparison, there was significant enrichment in amplification of the SE overlapping windows compared to the random groups (Figures 2D and E). This observation was reinforced by comparing SE CNV to the CNV across the ovarian cancer genome as a whole (Figure 2F). Remarkably, amplification of the super-enhancers themselves was prognostic of clinical outcome^39^. In many cases, as seen for the super-enhancer on chromosome 20 (chr20:55890001-55905000), patients with increased copy number had significantly increased hazard ratio and reduced survival times, suggesting that super-enhancer copy number may be of prognostic value (Figure 2C). Taken together, these data suggest that the SEs we identified in OVCAR3 cells are preferentially amplified in ovarian cancer patients and that some SE amplifications are associated with reduced survival.

**Figure 2.**
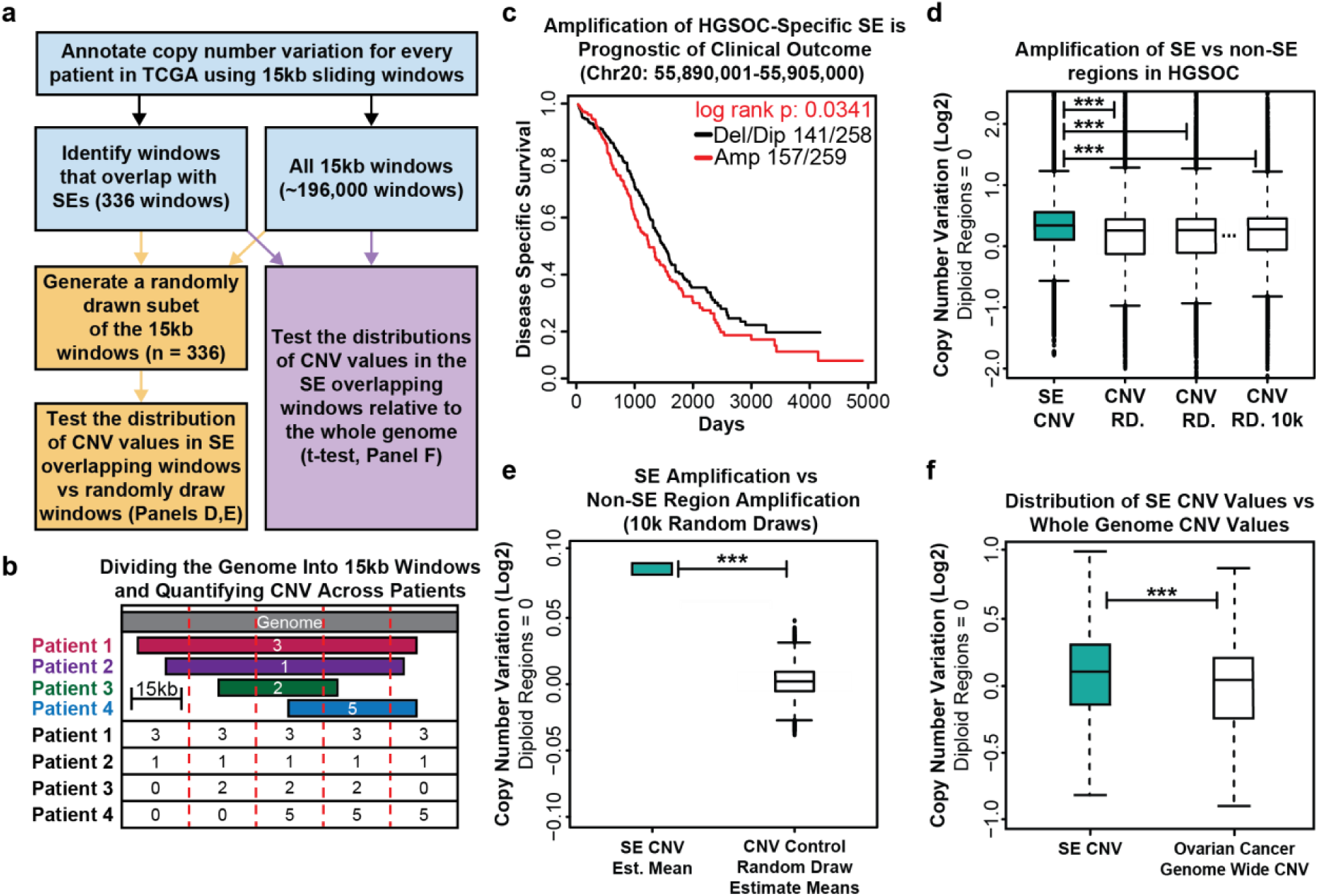
BRD4 Bound Super-Enhancers are Enriched for Copy Number Alterations in Ovarian Cancer Patients. **a.** Flowchart of the analysis strategy used to quantify the relationship between SEs defined in OVCAR3 cells and copy number alterations in high grade serous ovarian cancer patients. **b.** Cartoon showing the computational approach used to divide the genome into 15kb windows and assign patient-specific copy number values to each window by overlap analysis. **c.** Copy number Kaplan Meier plot for a 15kb window that overlaps an OVCAR3 defined SE at Chr20 55890001:55905000. The red line represents HGSOC patients with copy number amplification of this region above the median, the black line represents patients with copy number below the median. **d.** Boxplots showing the comparison of HGSOC patient copy number across SE overlapping windows (n = 336) versus randomly drawn genomic windows of the same size (n = 336). Asterisks represent significant differences as determined by a t-test. **e.** Summary plot showing the results of 10,000 comparisons between the copy number amplifications at SE overlapping windows versus 10,000 randomly drawn subsets of the genomic background. Asterisks represent significant differences as determined by a t-test. **f.** Boxplot showing the comparison of copy number amplification across the SE overlapping windows (n=336) versus all 15kb windows across the ovarian cancer genome (n=∼192,000). Asterisks represent significant differences as determined by a t-test.

To better understand how amplification of these SEs is associated with oncogenic gene expression networks, we leveraged the RNA-seq data generated from a subset of the same ovarian cancer patients (∼300) to link the SEs to gene expression. We took inspiration from a commonly used approach in complex genetics which associates nucleotide variants to changes in gene expression called eQTL analysis^40, 41^. However, unlike eQTL analysis which focuses on point mutations, the comparison in this case focuses on changes in copy number across SE loci to changes in gene expression within each patient (copy number variation expression quantitative trait loci (CNVeQTL)) (Supplemental Figure 2A). The assumption is that amplification or deletion of SE regions should affect their target genes, therefore, looking across hundreds of patients for shared patterns of variation will identify putative target genes of each SE. However, since we altered the input data of the eQTL detection software to utilize two quantitative variables (copy number and gene expression), we needed to determine a robust indication of our null condition for statistical analysis.

To generate the null dataset, we broke the linkage of RNA to copy number by randomly permuting the columns of the RNA data matrix and then running Matrix eQTL^41^ on this permutated dataset, repeating this process 100k times, and using the median distribution from all 100k trials to inform our experimental analysis. Importantly, all 100k runs using the permutated null data showed a relatively uniform distribution of *p*-values across the null condition, suggesting no meaningful relationship between copy number and gene expression, and had a similar count of total significant CNVeQTLs (the median number of CNVeQTLs across all 100k was 11,632) (Supplemental Figure 2B). In contrast, the results from the true data show a much sharper peak around *p*-value = 0 and returned a much larger number of significant CNVeQTLs (n=126,438) (Supplemental Figure 2C, Supplemental Data 5). We used the results of the 100k null experiments to determine an empirical false discovery rate of about 0.092^42^. This data also allowed us to investigate some higher order questions, such as whether the number of CNVeQTL detected was strictly a function of size. While there was a modest linear relationship between these features, this analysis suggested something other than genomic size influenced the number of CNVeQTL (Supplemental Figure 2D). Taken collectively, these data suggest that amplification of the super-enhancer regions are associated with pervasive gene expression changes in human tumors, reinforcing the idea they are not merely cell-line specific, and they may be preferentially amplified for a biologically meaningful reason.

We recognize that the identification of 126,438 CNVeQTL linkages across 126 super-enhancers seems high, despite the null distributions tested, and that the vast majority of copy number amplifications will have very strong effects in *cis* (and most will have effects in *trans*) irrespective of their designation as a super-enhancer. Therefore, to functionally validate and assess the full scope of this data, we chose the top 86 super-enhancers ranked by BRD4 enrichment and H3K27ac signal (which were located both above and below the CNVeQTL prediction line) to perturb using a high throughput CRISPRi screen (Supplemental Figure 2D).

### High Throughput CRISPR-Interference Screen Highlights Super-Enhancer Target Gene Relationships

To systematically probe the functions of each SE and determine the consequences on gene expression, we used high-throughput CRISPR-interference assays coupled to RNA-seq. For this experiment, we engineered OVCAR3 cells to stably express nuclease deficient Cas9 fused to the KRAB effector domain (dCas9-KRAB). The KRAB effector domain induces local chromatin repression via methylation of histone 3 lysine 9 (H3K9me3) and, when fused to dCas9, allows us to use the programmable properties of CRISPR to target and inhibit any genomic loci of interest (Figure 3A)^43–46^. For this experiment, each well received a different set of custom designed guide RNAs (sgRNAs) to specifically inhibit one SE per well (i.e. arrayed CRISPRi screen) (Figure 3B, Supplemental Data 1). A total of 86 super-enhancers were tested plus 10 control wells. Two different sgRNAs, targeting the two highest BRD4 peak summits within each super-enhancer, were designed for each SE (*see Methods*)^47^. For negative controls, we used a non-targeting scrambled sgRNA in addition to an sgRNA designed to target a dormant region of the genome (Supplemental Data 1). Each sgRNA was cloned into the pX-sgRNA-eGFP-MI plasmid and transfected into its corresponding wells. After 72 hours of epigenetic silencing, RNA was purified from each well and barcoded to specifically track which super-enhancer was probed per well (96 total barcodes). The RNA was prepped and sequenced on an Illumina platform to measure changes in gene expression as a consequence of super-enhancer inhibition (Figure 3B). Given our intent to survey as many super-enhancers as possible and to have a better opportunity to find those that exhibited the most profound effects on gene expression, we decided to probe each SE once within the 96-well setup, prioritizing breadth over the inclusion of replicates (Figure 3B). Therefore, a traditional differential gene expression analysis pipeline (requiring the use of replicates) had to be eschewed in favor of something better able to handle our experimental setup. We took inspiration from previous analyses performed on large-scale perturbation databases, such as Connectivity Map project (CMap),^48^ and chose to focus on relative changes in rank for each gene (uprank or downrank) rather than traditional differential gene expression analysis or absolute expression counts. The resulting changes in rank could then be investigated across the entire dataset by iterating through a series of rank change cutoffs, identifying super-enhancers that affected significantly more genes at a particular cutoff as compared to the negative control wells (based on an empirical false discovery rate of 0.1) (*see Methods*). Any genes detected at these thresholds could then be tentatively assigned as target genes to each SE (Figure 3, Supplemental Data 6). To investigate whether a traditional *relative expression* approach would have identified similar target genes, we determined the log2 fold change of every gene for each SE relative to the controls. We then assess the relationship between gene expression determined by *relative change in rank* and *relative expression* for each SE. In every case, the correlation between log2 fold change (LFC) and rank change (RC) was highest when comparing each SE to itself, as opposed to all other SEs in the screen, suggesting that differential gene expression calculated in both ways gave similar results (Supplemental Figure 3A). Notably, some of the correlations were much stronger than others, leading us to focus on SEs with a LFC vs RC correlation value above the mean. Of particular interest was super-enhancer 14 (SE14) which exhibited a LFC vs RC correlation value of 0.95 (the highest in the entire dataset), suggesting particularly robust results for this SE (Supplemental Figure 3B). Confident that our rank change approach was adequately supported by this comparison we proceeded to look for SEs that exhibited the most profound effects on gene expression.

**Figure 3.**
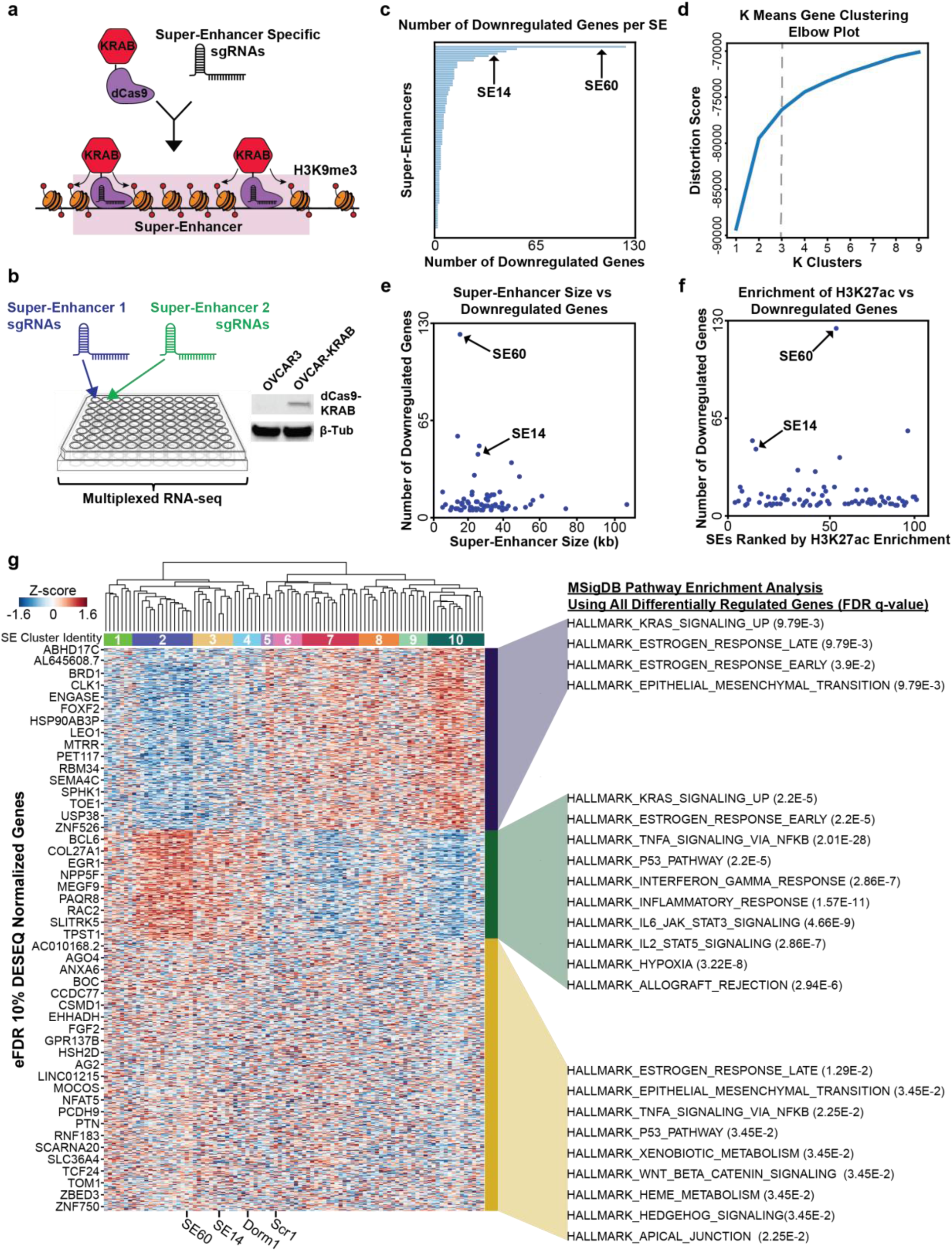
Systematic Epigenetic Silencing of Ovarian Cancer Specific Super-Enhancers using CRISPRi (dCas9-KRAB) Coupled to Multiplexed RNA-seq. **a.** Cartoon showing sgRNA guided dCas9-KRAB epigenetic silencing of a SE via enrichment of the repressive histone modification H3K9me3. **b.** Experimental setup for the CRISPRi screen in a 96-well plate (*left*). Western blot showing OVCAR3 cells engineered to stably express dCas9-KRAB (*right*). dCas9-KRAB expressing OVCAR3 cells were plated in each well. One SE was targeted per well (86 SEs plus 10 control wells). After 72 hours of enhancer silencing, changes in gene expression were measured using barcoded RNA-seq. Two sgRNAs were custom designed for each SE and transfected into each corresponding well. **c.** Horizontal bar chart showing the number of downregulated genes for each SE. SE60 and SE14 were selected for further analysis as described in the text and are indicated by arrows. **d.** K means clustering elbow plot used to determine the optimal number of gene clusters across significant DEGs for all SEs pulled from the screen analysis. The “elbow” determines the ideal cluster number which was chosen as 3. **e.** Scatterplot comparing SE size versus number of downregulated genes. There is no correlation between SE size and the number of target genes. SE60 and SE14 were selected for further analysis as described in the text and are indicated by arrows. **f.** Scatterplot comparing H3K27ac enrichment versus number of downregulated genes. There is not a strong correlation between H3K27ac enrichment and number of target genes. **g.** Heatmap representing the unsupervised hierarchical clustering of all SEs (clusters 1-10 under the dendrogram) and controls in the screen across all screen DEGs (*left*). The boxes on the right denote the three K-means clusters. MSigDB pathway analysis describes the functions the genes in these clusters are involved in (*right*). SE60, SE14, and the two negative controls are denoted at the bottom of the plot.

First, we focused on a number of summary analyses from the CRISPRi screen. The median number of genes downregulated by each SE was 4 and there were a few salient SEs that affected a much larger number of genes (Figure 3C). Interestingly, there was only a weak correlation between the number of differentially regulated genes and SE size (Figure 3E) or enrichment of H3K27ac (Figure 3F), suggesting that the effects on gene expression are not merely a function of size. Of note, super-enhancer 60 (SE60) was in the bottom half in terms of size, but it affected the greatest number of genes. Therefore, we felt it prudent to understand specificity of our CRISPRi targeting process and empirically determine the extent of spreading of the repressive H3K9me3 mark upon dCas9-KRAB binding. To this end, we performed H3K9me3 ChIP-seq in ovarian cancer cells transfected with SE60 targeting sgRNAs versus non-targeting sgRNAs. Differential binding analysis revealed that only our region of interest (SE60) was significantly enriched for H3K9me3 signal upon transfection of the targeting sgRNAs but not the scrambled non-targeting sgRNA (Supplemental Figure 4C and D). Additionally, there was an increase in H3K9me3 signal at each of our two SE60 sgRNA locations, suggesting both guides successfully delivered dCas9-KRAB to the SE (Supplemental Figure 4A and B). We found that the region with increased H3K9me3 was about 20kb, spreading ∼10kb from each sgRNA target site, enough to cover the entire SE. There also did not appear to be an increase in signal at the computationally predicted off-target sites, suggesting the guides for SE60 were highly specific (Supplemental Figure 4E and F). Taken together, these results validate our method of designing sgRNAs and the results we detected from the screen (specifically for SE60) were not due to off-target effects (Supplemental Figure 4).

Having supported the validity of our CRISPRi assay, we next wanted to examine the patterns of gene expression that resulted from the screen. To accomplish this, we utilized two clustering methods, K-Means clustering for the target genes and unsupervised-hierarchical clustering for the SEs. We found that the differentially regulated genes could be divided into 3 optimal clusters that represent distinct gene expression pathways in cancer cells (Figure 3D and G). Conversely, the SEs can be divided into 10 distinct clusters with shared patterns of gene expression (Supplemental Table 1). More specifically, CRISPRi targeting of the SEs in clusters 2-4 (containing SE14 and SE60) caused decreases in the expression of genes enriched for pathways such as *KRAS Signaling*, *Estrogen Response* (both early and late), and *Epithelial to Mesenchymal Transition (EMT)*. In contrast, SEs in clusters 5-10 maintain some similarities (*KRAS* and *Early Estrogen Response*) but also have a unique role in the regulation of the *JAK-STAT pathway* and immune related pathways (Figure 3G). Taken together, the CRISPRi screen in conjunction with our CNV analyses have allowed us to comprehensively determine which SEs have the most profound effects on gene expression and inform us of the enhancers that likely regulate key gene pathways in ovarian cancer. Based on these results, two salient SEs, SE60 and SE14, were selected for follow up experiments.

### Deletion of SE60 and SE14 Causes Dysregulation of Oncogenic Gene Expression Pathways Leading to Reduced Proliferation and Migration of Cancer Cells

Perturbation of SE60 affected the greatest number genes in the CRISPRi screen. In addition, amplification of this SE in ovarian cancer patients is prognostic of worse patient outcome, nominating it as a *bona fide* oncogenic super-enhancer (Figure 4A and B). Therefore, we wanted to experimentally determine whether SE60 was truly oncogenic and understand its role in ovarian cancer. To that end, we designed sgRNAs flanking the BRD4 peak summit of the largest constituent enhancer within SE60 and generated three independent CRISPR-Knockout (KO) clones resulting from ∼1700-1800bp deletions (Figure 4C).

**Figure 4.**
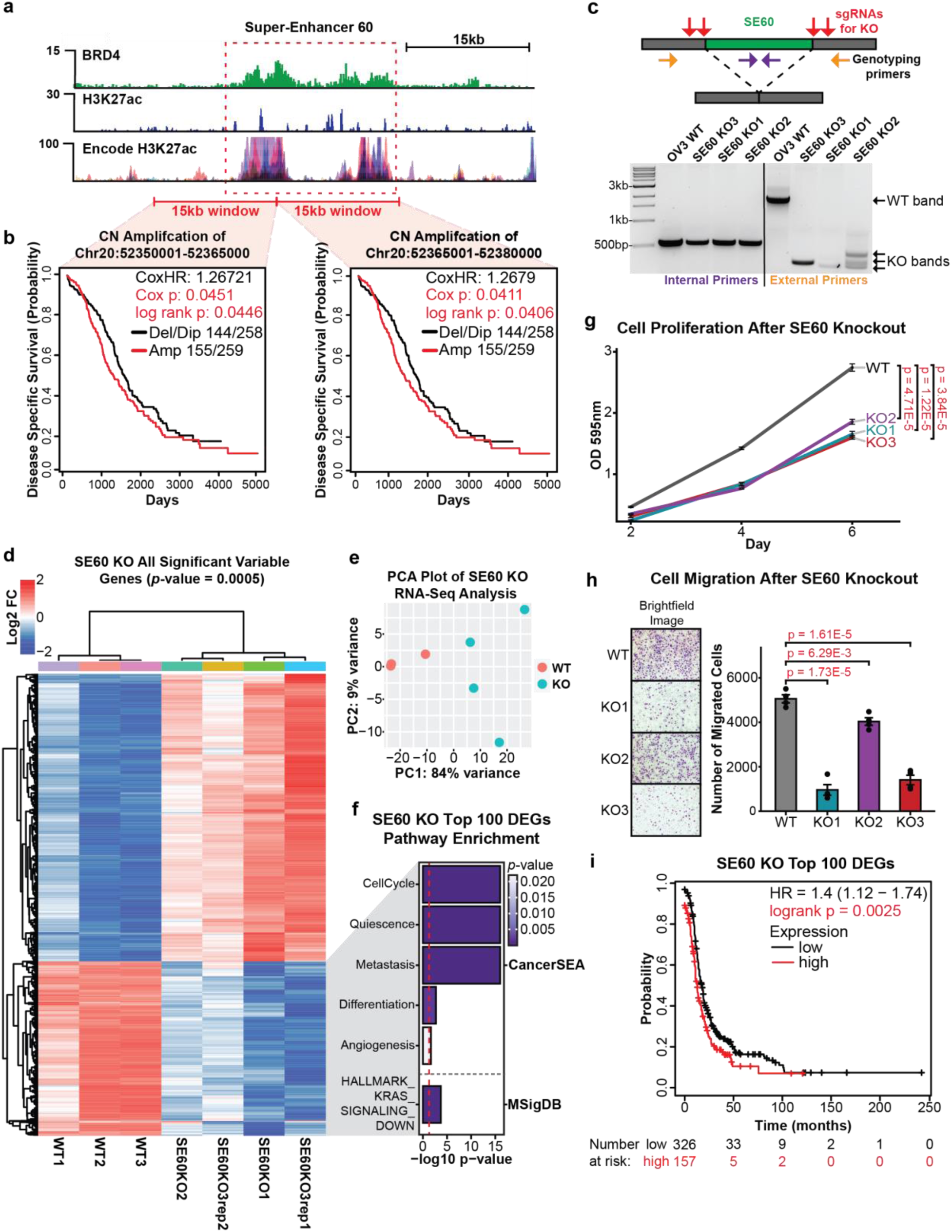
CRISPR-Knockout of Super-Enhancer 60 Leads to Profound Changes in Gene Expression and Reduced Proliferation of Cancer Cells. **a.** Genome browser view of SE60 (dashed red box) and the surrounding regions showing enrichment of BRD4, H3K27ac, and ENCODE H3K27ac signal. **b.** Copy number amplification of SE60 is prognostic of clinical outcome in HGSOC patients. Kaplan Meier plots of copy number amplification over each SE60 overlapping 15kb windows versus disease specific survival in TCGA HGSOC patients. Significance was assessed using a log-rank test and cox proportional hazards model. **c.** *Top:* Cartoon showing CRISPR-based deletion of SE60. *Bottom:* Genotyping PCR agarose gel electrophoresis showing successful heterozygous knockouts of SE60. Homozygous deletions were lethal to the cells and heterozygotes were severely impaired (see proliferation assays in *panel g and* invasion assays in *panel h*). **d.** *Left:* Unsupervised hierarchical clustering heatmap of all 1750 significant DEGs (adjusted p-value > 0.0005 at any fold change) between wild-type and SE60 KO cells measured by RNA-seq. **e.** PCA plot showing the variance landscape of all 3 WT and all 4 KO samples. **f.** Pathway analysis using CancerSEA and MSigDB of the 100 most significant DEGs detected in the analysis; the red line denotes the metric for a *p*-value of 0.05 (significance) converted into the -log10 scale, indicating significant terms. **g.** Proliferation assays of three independent KO clones (represented in the RNA-Seq data) of SE60 versus wild-type OVCAR3 cells. The statistically significant differences (as determined by a Welch’s t-test) are provided in red text. **h.** Cell Migration assays of three independent KO clones of SE60 (represented in the RNA-Seq data) versus wild-type OVCAR3 cells. Microscope brightfield images of the growth after 24 hours (*left*). Bar chart representation of cell count after 24 hours, statistically significant differences (as determined by a Welch’s t-test) are provided in red text (*right*). **i.** Kaplan Meier plot showing the clinical significance of the top 100 downregulated genes (as a gene set signature) after SE60 KO, the red line denotes patients with high expression of this gene signature. Significance was assessed using a log-rank test, significant *p*-values are denoted in red text.

Global changes in gene expression resulting from each SE60 KO clone were measured using RNA-seq. Differential expression analysis using DESeq2^49^ revealed pervasive changes in gene expression with 660 genes being detected as significantly downregulated and 1090 genes being upregulated at a strict confidence threshold (adjusted *p*-value of 0.0005) (Figure 4D and E, Supplemental Data 7). Pathway analysis of the top 100 significantly downregulated genes

(determined by *p*-value) identified significant enrichment in *Cell Cycle Progression*, *Quiescence*, *Metastasis*, *Differentiation*, and *KRAS-Signaling* (Figure 4F), further suggesting SE60 plays an important role in oncogenesis^50^. This observation was supported upon clinical analysis of these predicted target genes, where increased expression of the top 100 SE60 target genes is associated with worse clinical outcomes in HGSOC patients (Figure 4I). The notion that SE60 plays a key role in ovarian cancer oncogenic processes was further validated by the profound effects that deletion of this SE had on cancer cell proliferation and migration (Figure 4G and H).

To substantiate our approach for identifying clinically relevant oncogenic super-enhancers, we selected an additional candidate from the CRISPRi screen for validation. SE14 showed the highest correlation between LFC and RC differential gene expression analysis from the CRISPRi screen, it is in the top 4 SEs that affected the greatest number of genes, and its amplification portends a worse clinical outcome in ovarian cancer patients (Figure 5A and B). To investigate the functional role of SE14, we designed sgRNAs flanking the BRD4 peak summit of the largest constituent enhancer within the super-enhancer and generated three independent CRISPR-KO clones resulting from ∼2500-2800bp deletions (Figure 5C). Global changes in gene expression resulting from each knockout clone were measured using RNA-seq. Differential expression analysis identified 860 genes as significantly downregulated, and 629 genes upregulated at our confidence threshold (adjusted *p*-value of 0.0005) (Figure 5D and E, Supplemental Data 7). Pathway analysis of the top 100 most significant downregulated genes identified significant enrichment in *Cell Cycle Progression*, *Quiescence*, *Metastasis*, *Differentiation*, and *EMT* (Figure 5F), further suggesting SE14 plays an important role in oncogenesis^50^. Kaplan-Meier analysis of the top 100 most significant downregulated genes after SE14 KO revealed a significant association with worse clinical outcomes in HGSOC patients (Figure 5I). Similar to the results obtained with SE60, the biological assays on all three SE14 KO cell lines exhibited a significant decrease in proliferation and migration compared to wild type cells (Figure 5G and H).

**Figure 5.**
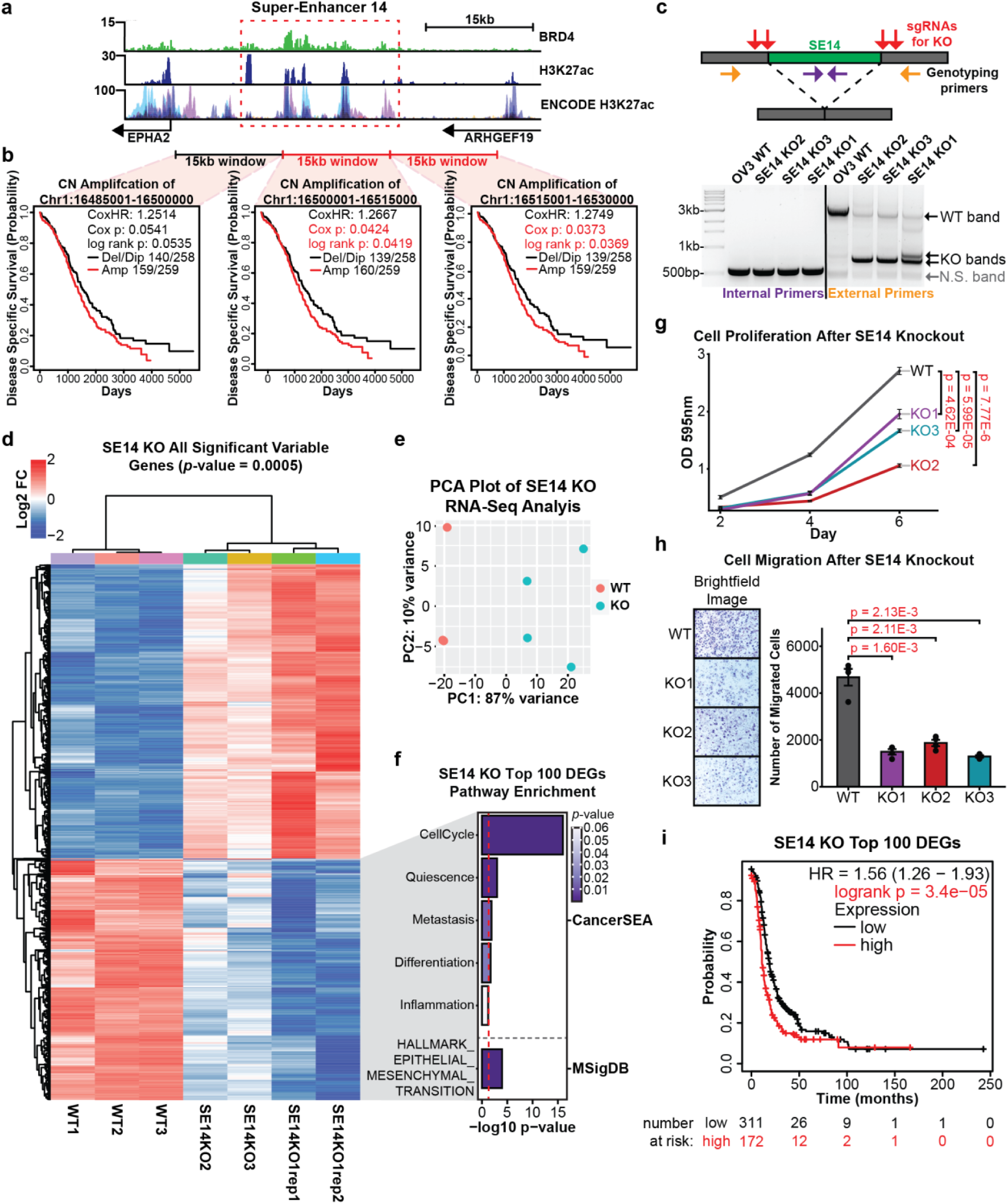
CRISPR-KO of Super-Enhancer 14 Leads to Important Gene Expression Changes and Reduced Growth of Cancer Cells. **a.** Genome browser view of SE14 (dashed red box) and the surrounding regions showing enrichment of BRD4, H3K27ac, and ENCODE H3K27ac signal. **b.** Copy number amplification of SE14 is prognostic of clinical outcome in HGSOC patients. Kaplan Meier plots of copy number amplification over each SE14 overlapping 15kb windows versus disease specific survival in TCGA HGSOC patients. Significance was assessed using a log-rank test and cox proportional hazards model. **c.** *Top:* Cartoon showing CRISPR-based deletion of SE14. *Bottom:* Genotyping PCR agarose gel electrophoresis showing successful heterozygous knockouts of SE14. Homozygous deletions were lethal to the cells and heterozygotes were severely impaired (see proliferation assays in *panel g and* invasion assays in *panel h*). **d.** *Left:* Unsupervised hierarchical clustering heatmap of all 1750 significant DEGs (adjusted *p*-value > 0.0005 at any fold change) between wild-type and SE14 KO cells measured by RNA-seq. **e.** PCA plot showing the variance landscape of all 3 WT and all 4 KO samples. **f.** Pathway analysis using CancerSEA and MSigDB of the 100 most significant DEGs detected in the analysis; the red line denotes the metric for a *p*-value of 0.05 (significance) converted into the -log10 scale, indicating significant terms. **g.** Proliferation assays of three independent KO clones (represented in the RNA-Seq data) of SE14 versus wild-type OVCAR3 cells. The statistically significant differences (as determined by a Welch’s t-test) are provided in red text. **h.** Cell Migration assays of three independent KO clones of SE14 (represented in the RNA-Seq data) versus wild-type OVCAR3 cells. Microscope brightfield images of the growth after 24 hours (*left*). Bar chart representation of cell count after 24 hours, statistically significant differences (as determined by a Welch’s t-test) are provided in red text (*right*). **i.** Kaplan Meier plot showing the clinical significance (hazard) of the top 100 downregulated genes (as a gene set signature) after SE60 KO, the red line denotes patients with high expression of this gene signature, significant *p*-values are denoted in red text.

We had an interest in determining how similar the results of CRISPRi-based perturbation of SE60 and SE14 are to the gene expression changes caused by CRISPR-KO. Therefore, we performed additional dCas9-KRAB experiments coupled to RNA-seq (in replicate) for both SE60 and SE14. Differential gene expression analysis for both the CRISPRi and CRISPR-KO was performed with DESeq2 to facilitate the comparisons of the resulting changes in gene expression (Supplemental Figure 5, Supplemental Figure 6, and Supplemental Data 7). For SE60, 169 genes were detected as differentially expressed by both CRISPRi and CRISPR-KO and 11 of these genes were downregulated, suggesting that these are true target genes of SE60 (Supplemental Figure 5C and D). Further analysis of the 11 downregulated genes detected by both CRISPRi and CRISPR-KO found this gene set to be enriched for *Metastasis*, *Cell Cycle Progression*, and *Inflammation* pathways, as well as being associated with reduced survivorship in HGSOC patients (Supplemental Figure 5F and 5G). The analysis of SE14 revealed 731 differentially expressed genes by both CRISPRi and CRISPR-KO and 169 of these genes being downregulated (Supplemental Figure 6C and D). Analysis of the 169 shared downregulated genes detected by both CRISPRi and CRISPR-KO found this gene set to be enriched for *Quiescence*, *Cell Cycle Progression*, *Differentiation*, *Inflammation*, *Stemness*, and *Estrogen Response* pathways as well as being associated with reduced survivorship in HGSOC patients, further reinforcing the role of SE14 in oncogenesis (Supplemental Figure 6F and 6G). Taken together, these results validate our oncogenic SE identification approach and highlight the importance of these two SEs in ovarian cancer. Next, we investigated whether the mechanistic roles of SE60 and SE14 on cell proliferation and migration were due to direct or indirect target gene regulation.

### 3D-Chromatin Interactions Defined by Hi-C Sequencing of Ovarian Cancer Cells Establish Direct Target Genes for SE60 and SE14

The profound effects on proliferation and migration caused by CRISPR-based deletion of SE60 and SE14 led us to investigate if these biological phenotypes were caused by *direct* or *indirect* regulation of target genes. We reasoned that direct target genes would exhibit increased chromatin looping interactions with the SE, whereas indirect target genes would be downstream of an effector gene that was directly regulated by the SE. To enable unbiased measurement of interaction frequencies between each super-enhancer and its target genes, we performed Hi-C sequencing in OVCAR3 cells to comprehensively annotate chromatin interactions across the ovarian cancer genome^51^. In order to maximize the breadth of this analysis, we focused on the target gene set detected from the CRISPR-KO experiments that represented the most statistically robust gene set for each SE, resulting from 4 replicates of RNA-seq across three independent knockout clones for each SE. Moreover, the constitutive perturbation of each SE, caused by CRISPR-based deletion, gave rise to consistent gene expression patterns that resulted in marked biological phenotypes, thus facilitating integration with the Hi-C data.

Since Hi-C is highly dependent on distance, we limited our search space to genes located on the same chromosome as the super-enhancers (*cis* genes) in order to get an accurate metric of interaction frequency^52, 53^. To perform this analysis, we compared interaction frequency between the SE and target gene pairs to a null distribution consisting of 100 permutations of distance-matched region-gene pairs that exhibited no significant changes in gene expression changes in gene expression upon SE deletion (*see Methods*). This enabled us to compare distributions of interaction frequency measurements between each SE and a random set of genes based entirely on genomic distance. Direct targets were defined as SE-gene pairs with an observed/expected contact frequency greater than the 75th percentile of the control/background distribution. Overall, we observed that the target genes for each SE had higher interaction frequency with their cognate SE as compared to an equivalent number of distance-matched genes found on the same chromosome (Figure 6A and F).

**Figure 6.**
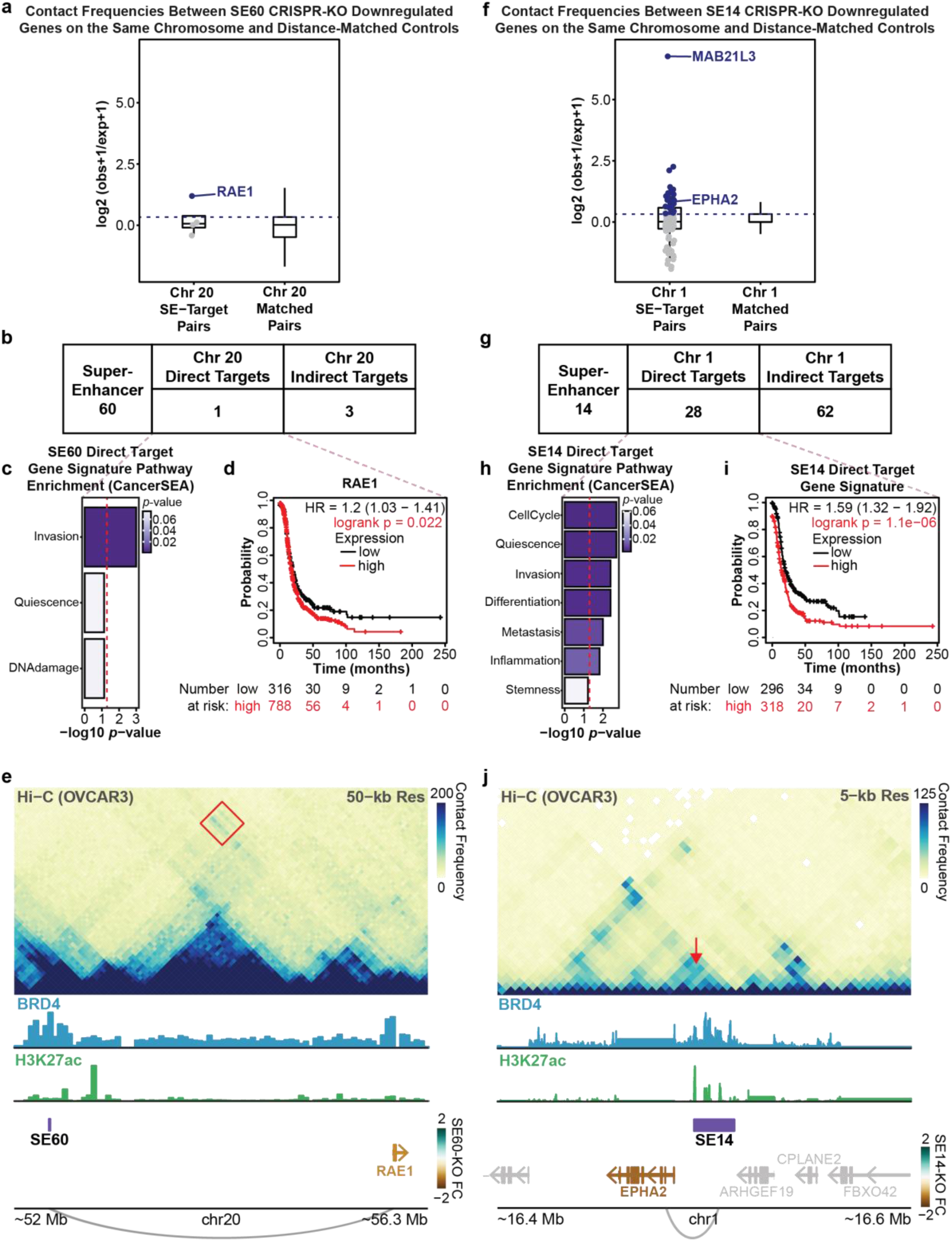
Hi-C Analysis Detects Direct Targets of SE60 and SE14 Supporting Direct Roles in Invasion, Differentiation, and Metastasis. **a.** Distribution of all Hi-C counts (contact frequency) between cis-gene SE60 downregulated DEGs and the SE60 locus (*left*), points/genes in blue are called as direct targets. Distribution of Hi-C counts (contact frequency) between SE60 and a background set comprised of 100 distance permuted control gene sets (*right*). The dashed line (3^rd^ quantile of the background data) denotes the cutoff for direct target genes in the experimental sample. **b.** Table displaying the number of direct and indirect cis-target genes of SE60 determined from the Hi-C analysis. **c.** CancerSEA gene pathway analysis of the direct target genes detected in the analysis; the red line denotes the metric for a *p*-value of 0.05 (significance) converted into the -log10 scale, anything past the line is a significant term. **d.** Kaplan Meier plot showing the clinical significance (hazard) of the top direct target genes (as a gene set signature) after SE60 KO, the red line denotes patients with high expression of this gene signature, significant *p*-values are denoted in red text. **e.** Hi-C contact heatmap showing the interaction between *RAE1*, the direct target gene of SE60, and the SE locus itself (*red square*). **f.** Distribution of all Hi-C counts (contact frequency) between cis-gene SE14 downregulated DEGs and the SE14 locus (*left*), points/genes in blue are called as direct targets. Distribution of Hi-C counts (contact frequency) between SE14 and a background set comprised of 100 distance permuted control gene sets (*right*); the dashed line (3^rd^ quantile of the background data) denotes the cutoff for direct target genes in the experimental sample. **g.** Table displaying the number of direct and indirect cis-target genes of SE14 determined from the Hi-C analysis **h.** CancerSEA gene pathway analysis of the direct target genes detected in the analysis; the red line denotes the metric for a *p*-value of 0.05 (significance) converted into the -log10 scale, anything past the line is a significant term. **i.** Kaplan Meier plot showing the clinical significance (hazard) of the top direct target genes (as a gene set signature) after SE14 KO, the red line denotes patients with high expression of this gene signature, significant *p*-values are denoted in red text. **j.** Hi-C contact heatmap showing the interaction between *EPHA2*, a direct target gene of SE14, and the SE locus itself (*red arrow*).

We identified one *cis* direct target gene and four *cis* indirect target genes for SE60 (Figure 6B). Of note, the *cis* direct target gene for SE60, *RAE1,* has previously been associated with *Invasion* in ovarian cancer and has been shown to promote *EMT* in breast cancer^54^. In addition, increased expression of this gene portends a worse outcome in HGSOC patients (Figure 6C and D). Notably, *RAE1* was also predicted as a target of SE60 by the CNVeQTL analysis (Supplemental Figure 8A and B, Supplemental Data 8). When looking at Hi-C contact frequency across chromosome 20, we notice a marked increase in contact between the *RAE1* locus and the SE60 locus as compared to the background (Figure 6E). This suggests that the decrease in migration detected upon SE60 deletion is due, in part, to its direct regulation of *RAE1*. We suspect that there may exist more direct target genes for SE60 located on other chromosomes whose interaction frequencies are technically more challenging to detect via Hi-C.

Interestingly, we identified a much greater number of *cis* direct target genes (28 genes) and *cis* indirect targets (62 genes) for SE14 (Figure 6G, Supplemental Data 8). Likewise, 8 of these *cis* direct targets had been predicted from our CNVeQTL analysis reinforcing the utility of CNVeQTLs to predict *cis*-direct targets (Supplemental Figure 8C and D). Pathway analysis of the *cis* direct targets revealed key roles in *Cell Cycle Progression*, *Quiescence*, *Invasion*, *Differentiation*, Metastasis, and *Stemness* (Figure 6H). Kaplan-Meier analysis of this gene signature highlighted a statistically significant decrease in survival for patients that had high expression of these genes (Figure 6I). Through our analysis of these of *cis* direct targets, we identified examples of both close-range (*EPHA2*) and distant (*MAB21L3*) connections to SE14 (Figure 6J and Supplemental Figure 7). Interestingly, there were genes within very close proximity to SE14 (such as *ARHGEF19)* that showed no evidence of interaction or differential gene expression, highlighting the ability of our approach to delineate true targets when accounting for distance. These data strongly implicate SE14 as being directly involved in both proliferation and migration as well as other key oncogenic processes in ovarian cancer.

### SE60 and SE14 are Specifically Active Within the Epithelial Cancer Cell Fraction of Human HGSOC Tumors as Revealed by Single Cell Genomics

Our previous experiments had demonstrated that these SEs are preferentially amplified in ovarian cancer patients and that they regulate gene expression pathways that govern the proliferation of cancer cells. As a final validation experiment, we wanted to determine if SE60 and SE14 were specifically active within the cancer cell compartment of human HGSOC tumors and if their target genes are also active within the same cell type. To test this, we analyzed matched single-cell RNA-seq and single-cell ATAC-seq data from two HGSOC patients previously generated in our lab (Supplemental Figure 9)^38^. We annotated seven distinct cell types present in these tumors by both single-cell RNA-seq and single-cell ATAC-seq and identified the cancer cell population using the FDA approved biomarker CA125 (also known as *MUC16*) (Figure 7A and B)^55^. We found significant enrichment of *RAE1*, a SE60 *cis* direct target, and *EPHA2*, a SE14 *cis* direct target, within the cancer cell fraction as compared to the normal cell fraction (Wilcoxon Rank Sum tests, Bonferroni-corrected *p*-values < 2.2e-308 & average logFC >= 0.1) (Figure 7B). In order to assess whether SE60 and SE14 are uniquely active in ovarian cancer, we next leveraged the scATAC-seq data. These data showed significantly increased chromatin accessibility at three constituent enhancers of both SE60 and SE14, specifically within the cancer epithelial cell fraction as compared to the stromal compartments of these tumors (Wilcoxon Rank Sum tests, Benjamini-Hochberg FDR <= 0.10 & Log2FC >= 0.25). Additionally, both HGSOC patients showed this pattern, suggesting that activation of these SEs is a common feature of HGSOC biology. While there is previous evidence from ENCODE that these regions contain regulatory elements in normal ovarian tissue, it appears that there is significantly more accessibility of these super-enhancers in cancer cells (Figure 7C).

**Figure 7.**
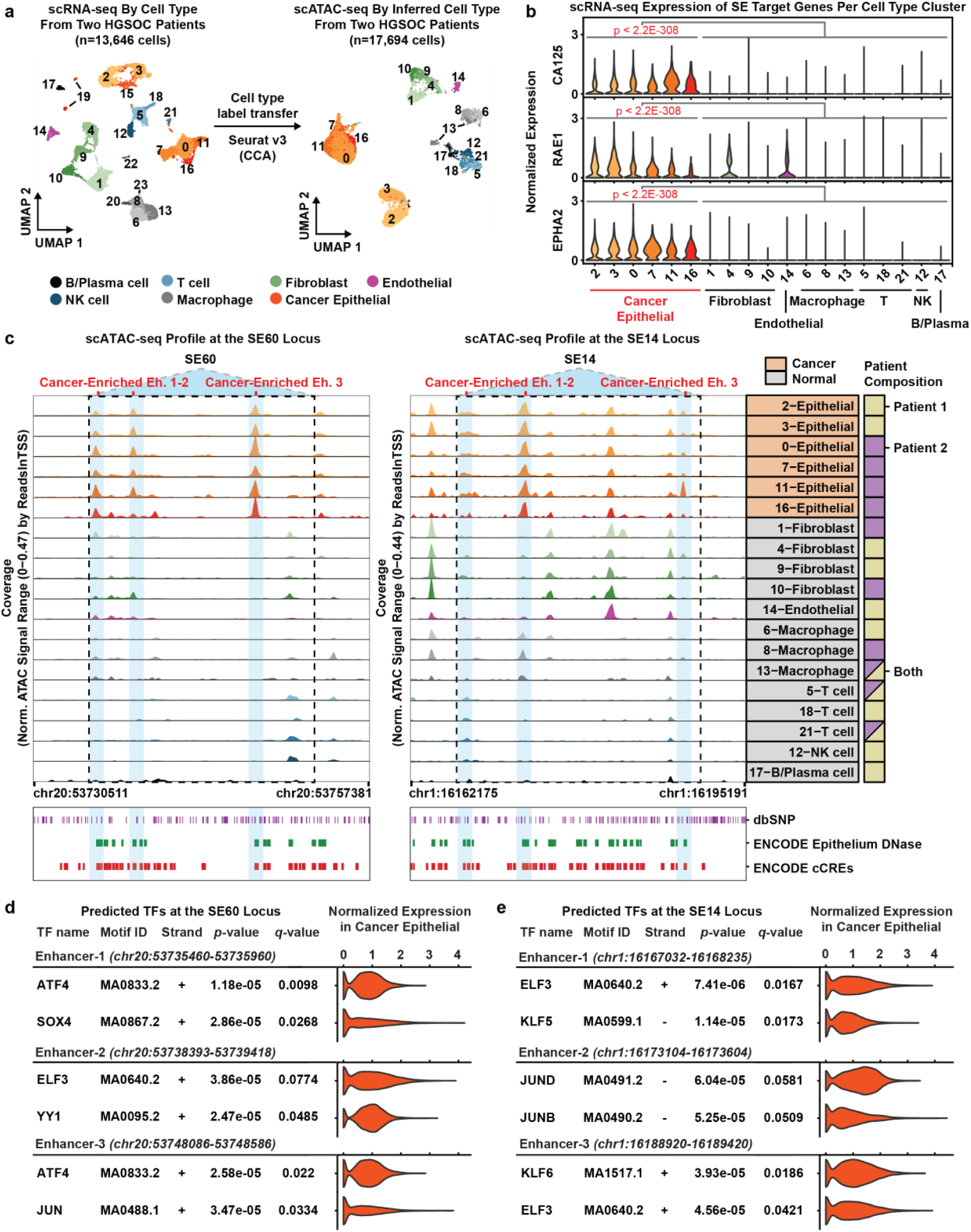
Super-Enhancer 60, 14, and Their Direct Target Genes are Enriched in Malignant Cells of HGSOC Patient Tumors as Determined by Single Cell RNA-seq and Matched Single Cell ATAC-seq. **a.** UMAP plot of 13,646 scRNA-seq cells colored by cell type from two HGSOC patient tumors (*left*). UMAP plot of 17,694 scATAC-seq cells from the same two HGSOC patient tumors colored by cell type (*right*). Cluster numbers in each UMAP plot denote cell type clusters. Only cell type clusters with at least 30 cells in scATAC-seq were used in downstream analyses and are labeled on the scATAC-seq UMAP plot (*right*). **b.** Violin plots showing the distribution of gene expression values measured by scRNA-seq in each cell type cluster. Each row shows the distributions of expression values for a single gene (*CA125*, *RAE1*, and *EPHA2*). Each column represents a cell type cluster denoted by a cluster number and a general cell type label (*bottom*). Each gene has a statistically significant difference in expression between the cancer and non-cancer cell type clusters (Wilcoxon Rank Sum tests, Bonferroni-corrected *p*-values < 2.2e-308 & average logFC >= 0.1). **c.** scATAC-seq browser track showing the chromatin accessibility profile at the SE60 locus for each cell type cluster in scATAC-seq (*left*). scATAC-seq browser track showing the chromatin accessibility profile at the SE14 locus for each cell type cluster in scATAC-seq (*right*). Light blue shadows denote cancer enriched constituent enhancer elements. Each light blue region has a statistically significant difference in accessibility between the cancer and non-cancer cell type clusters (Wilcoxon Rank Sum tests, Benjamini-Hochberg FDR <= 0.10 & Log2FC >= 0.25). Cancer status is denoted in each browser track row label where the cell type cluster is orange if the cells are from the cancer fraction of patients. Patient composition is denoted by the square to the right of the label, it is solid if it contains cells only from one patient or split colored if otherwise (far *right*). The dbSNP, Epithelium DNase, and ENCODE ccREs tracks denote the location of annotated SNPs, ENCODE DNase hypersensitivity sites in normal epithelium tissue samples, and ENCODE annotated regulatory elements, respectively (*bottom*). **d.** Summary of FIMO TF motif occurrences within SE60 cancer enriched enhancers 1-3. Matching scRNA-seq TF expression in the cancer epithelial fraction is shown in the violin plot for each predicted motif. Statistically significant motif matches identified by the FIMO software were defined as a Benjamini-Hochberg corrected *p*-value (i.e., q value) < 0.10. **e.** Summary of FIMO TF motif occurrences within SE14 cancer enriched enhancers 1-3. Matching scRNA-seq TF expression in the cancer epithelial fraction is shown in the violin plot for each predicted motif. Statistically significant motif matches identified by the FIMO software were defined as a Benjamini-Hochberg corrected *p*-value (i.e., q value) < 0.10.

In order to investigate what transcription factors might be involved with these super-enhancers, we performed motif enrichment analysis using FIMO sequence analysis^56^. To provide confidence to the TF motif calls, we investigated the gene expression of the TFs within the cancer epithelial cells. Transcription factors such as SOX4, ATF4 and YY1 were significantly enriched in the cancer-enriched constituent enhancers of SE60. Of note, YY1 is known as an integral component of enhancer-promoter loop interactions and is a hallmark active enhancer^57^. Similarly, in SE14 we detected significant enrichment of ELF3, KLF, and JUN family member binding sites in the cancer-enriched constituent enhancers. ELF3 has previously been previously associated with vascular inflammation, tumorigeneses, epithelial differentiation, and the ERRB3 pathway providing additional evidence to the robustness of our analysis (Figure 7D and E)^58, 59^. Taken together, these data suggest that these SEs and their target genes are cancer cell specific and validate our computational pipeline for identification of clinically relevant oncogenic super-enhancers.

## DISCUSSION

Every year, an estimated 22,000 new cases of ovarian cancer will be diagnosed and around 14,000 women will die as a result of this disease^4^. The paucity of known drivers for ovarian cancer makes identifying at-risk individuals very difficult and has led to a lack of effective targeted therapies. Thus, platinum-based chemotherapy coupled with surgery remains the standard of care^10^. Given their critical functions in controlling gene regulation, enhancers are often required to achieve the levels of transcriptional activity needed to sustain cancer cells and have been shown to play an integral part in cancer development and patient survival. Additionally, super-enhancers have demonstrated the capacity to regulate many critical pathways for the development and maintenance of the cancer cell state as well as influence therapeutic resistance^18–21^.

With the advent of therapeutics designed to inhibit various epigenetic factors that convey functionality to enhancers, it is now possible to exploit the dependency of cancer cells on transcription as an effective strategy for treating therapeutically recalcitrant cancers such as ovarian cancer^28, 60^. For example, the Bromodomain and Extra-Terminal motif inhibitors (BET inhibitors; such as JQ1) designed to interfere with the functions of bromodomain-containing proteins like BRD4 have shown promise in several pre-clinical models of cancer, although their efficacy in a clinical setting is still unknown^26, 61^. Nonetheless, investigating enhancers with high BRD4 enrichment can lead to the identification of biomarkers, druggable targets, and an improved understanding of ovarian cancer. Notably, the expression of BRD4 is highest in ovarian cancer as compared to every other cancer type represented in The Cancer Genome Atlas and high expression portends a worse outcome in ovarian cancer patients (Figure 1). Thus, we reasoned that co-enrichment of BRD4 and H3K27ac can be used as a surrogate to find SEs driving oncogenic processes in ovarian cancer. This was substantiated by the observation that the SEs identified in our study were preferentially copy number amplified in ovarian cancer patients and that some amplification events were themselves predictive of worse clinical outcome (Figure 2). Additionally, our CNVeQTL analyses across HGSOC patients demonstrate that the activity of these super-enhancers is pervasive. This is perhaps not surprising since genomic instability is a hallmark of ovarian cancer and several studies have demonstrated that somatic mutations at specific regulatory elements in the ovarian cancer genome play a pivotal role in subtype determination and overall progression^5, 62^. Furthermore, the dysregulation of genomic architecture in ovarian cancer may allow for cancer cells to hijack existing enhancers for oncogenic purposes. In fact, several examples of enhancer hijacking exist in other types of cancer such as in Burkett’s Lymphoma, B-Cell Lymphoma, and Glioblastoma^3, 63, 64^. Overall, these findings suggested that a number of our identified SEs were amplified for biologically meaningful reasons.

Rather than limiting our study to the standard taxonomic listing of super-enhancers, we used three orthogonal approaches to define the regulatory logic of SEs in ovarian cancer - CRISPRi, CRISPR-KO, and Hi-C. The CRISPRi screen allowed us to systematically determine the target genes for each of the top 86 most active SEs (Figure 3). While most CRISPR screens involve a pool of sgRNAs and rely on a cellular endpoint such as proliferation to be able to capture the relative abundances of remaining sgRNAs, our screen was customized to provide a readout of gene expression for each super-enhancer. We knew, *a priori*, which sgRNAs were used and which SEs were affected in each well. On average, we found that each SE perturbation resulted in downregulation of about 4 genes and the total number of genes regulated by each SE was not a function of size or enrichment of H3K27ac or BRD4. In fact, SE60 was in the bottom quartile of super-enhancers in terms of size and H3K27ac enrichment, but it had the most profound effects on gene expression. Therefore, we reasoned that SE60 harbored the most potential for further study due to its likely role in regulating genes that contribute to the pathology of ovarian cancer. While the goal of the CRISPRi screen was to broadly investigate the effects on gene expression across a large cohort of super-enhancers, we recognize that the CRISPRi screen was underpowered to definitively establish target genes for each SE. Thus, we elected to perform CRISPR-KOs of SE60 and SE14 to enable robust target gene detection.

CRISPR-KO of SE60 and SE14 had dramatic effects on gene expression programs involved in *Quiescence*, *Metastasis*, and *Invasion,* among other important pathways. Moreover, the genes sets for both SEs were associated with poor outcomes in HGSOC patients. This was supported by both proliferation and migration defects in the SE60 and SE14 knockout cells (Figure 4 and Figure 5). We note that there were hundreds of genes differentially regulated upon deletion of these two SEs, and that it was important to determine which genes were direct targets. The field has wrestled with the best way to assign target genes to enhancers, especially considering the genomic rearrangements observed in cancer cells. We reasoned that direct chromatin interactions, measured via Hi-C, between the SEs and their target genes would give confidence to the annotation of target genes (Figure 6).

Overall, the downregulated genes upon SE deletion showed higher interaction, both nearby and across long distances, with the SE as compared to the distance-matched control gene set. Importantly, several *cis* direct target genes are involved in oncogenic pathways and perhaps could serve as prognostic indicators or biomarkers in the future (Figure 6, Supplemental Figure 7, and Supplemental Figure 8). We do note that there may exist more direct target genes for each SE located on other chromosomes, however, due to the dependence of Hi-C on distance, these interaction frequencies are more technically challenging to quantify. In lieu of Hi-C data for determining direct target genes, we posit that evidence from two orthogonal experiments (such as CRISPRi and CRISPR-KO or inclusion of reporter-based enhancer assays) would yield high confidence results since genes detected by multiple assays are agnostic to the technical nuances of each. In fact, a logical framework to describe the level of support needed to definitively annotate an enhancer and its bona fide target genes has been recently proposed ^65^, and its implementation would yield a catalogue of enhancers with confidently linked target genes.

Finally, both SE60 and SE14 were found to have a statistically significant increase in chromatin accessibility within the cancer cell fraction of human HGSOC tumors at single cell resolution (Figure 7), further suggesting that the SEs that we identified are not merely cell-line specific. This validates our enhancer identification pipeline and reveals that certain oncogenic super-enhancers are preferentially enriched and amplified in cancer cells. In addition, we found that *cis* direct target genes annotated for each SE (such as *RAE1* and *EPHA2*) were more highly expressed in the cancer cells compared to the stromal/non-malignant cells within HGSOC tumors. Collectively, these results expound the concept that super-enhancers themselves and the genes they regulate represent viable therapeutic avenues and may aid in biomarker identification. More broadly, our study described a genomic and computational approach for identifying clinically relevant enhancers and their bona fide target genes which should be applicable to a wide variety of biological systems.

## METHODS

### Cell Culture

OVCAR3 and HEK-293T cell lines were obtained from ATCC. OVCAR3 cells were grown in RPMI media (Gibco, 11875-093) supplemented with 10% FBS (Sigma) and 1% penicillin/streptomycin (Corning, MT30002CI). HEK-293T cells were grown in Dulbecco’s Modified Eagle’s Medium (DMEM) (Gibco, 11995065) supplemented with 10% FBS and 1% penicillin/streptomycin. OVCAR3-dCas9-KRAB-blast (OVCAR-KRAB) cells were grown in RPMI media supplemented with 10% FBS, 1% penicillin/streptomycin and 1 µg/mL blasticidin (Corning, 30100RB) after selection. All cell cultures were incubated at 37 °C in 5% CO2. Before use, OVCAR3 cells were authenticated with Short Tandem Repeat profiling through ATCC. All cell lines were tested for mycoplasma.

### Engineering dCas9-KRAB expressing OVCAR3 cells

Lentivirus containing the Lenti-dCas9-KRAB-blast vector (Addgene plasmid #89567) was packaged in HEK-293T cells. HEK-293T cells were seeded in a T75 flask and transfected with the following plasmids: 6.67 µg Lenti-dCas9-KRAB-blast, 5 µg psPAX2 (Addgene, 12260), and 3.33 µg PMD2G (Addgene, 12259) using Fugene 6 (Promega, E2691) following the manufacturer’s protocol. The lentivirus containing supernatant was harvested 48-72 hours after transfection and lentivirus was concentrated using Lenti-X Concentrator (Takara, 631231) following the manufacturer’s protocol. OVCAR3 cells were seeded in a six-well plate at 50,000 cells/well and transduced with the harvested lentivirus in RPMI media with 10% FBS and 10 µg/mL polybrene (Millipore, TR1003G). Transduced cells were incubated with lentivirus for 72 hours, then placed in RPMI selection media with 3 µg/mL blasticidin for 7 days. Batch selected OVCAR3-KRAB cells were validated by western blot. For western blot analysis, cells were lysed using the following lysis buffer: 50 mM Tris HCl (pH 8), 0.5 M NaCl, 1% NP-40, 0.5% sodium deoxycholate, 0.1% SDS and 1x protease inhibitor. The primary antibodies used for Western blotting were as follows: β-tubulin (Abcam, ab6046), Cas9 (7A9-3A3) (Santa Cruz, sc-517386). The β-tubulin antibody was used at a 1:1500 dilution in 5% BSA in

TBST with overnight incubation at 4°C. The Cas9 antibody was used at a 1:1500 dilution in 5% BSA in TBST with overnight incubation at 4°C. The secondary antibodies used for Western blotting were as follows: Donkey anti-rabbit IgG HRP-linked (GE, NA934) and Donkey anti-mouse IgG HRP-linked (Invitrogen, PA1-28748). Secondary antibodies were used at a 1:5000 dilution in 5% BSA in TBST.

### CRISPRi Screen sgRNA Design

For sgRNAs targeting super-enhancers, target regions were chosen by selecting two regions within the super-enhancer with the highest BRD4 enrichment and clear H3K27ac signal. For each super-enhancer region, sgRNAs were designed using the CRISPOR web tool^47^ taking into account the specificity and off-target scores. If all suggested sgRNA sequences to a region had low specificity scores, a second sgRNA was instead designed to target the third highest BRD4 peak. Two sgRNAs were designed per super-enhancer to be transfected together. Genomic coordinates for all super-enhancers and their sgRNA sequences are found in Supplemental Data 1. sgRNA oligos were ordered from Integrated DNA Technologies. The negative control sgRNAs (Scramble1 and Scramble2) were previously published^66^.

### sgRNA Vector Cloning

The sgRNA cloning vector pX-sgRNA-eGFP-MI is a modified version of pSpCas9(BB)-2A-Puro (pX459) v2.0 (Addgene plasmid #62988). Cas9 was removed from pX459 and replaced with eGFP to allow for visualization of sgRNA expression. To improve sgRNA stability and optimize for assembly with dCas9, the sgRNA stem-loop was extended and modified with an A-U base pair flip^67^. sgRNA vector cloning was done following the protocol from Feng Zheng’s group^68^. Briefly, sgRNA oligonucleotides were ordered from Integrated DNA Technologies (IDT). Oligos were duplexed with the following reaction: 10 µM sgRNA forward oligo, 10 µM sgRNA reverse oligo, 10 U T4 polynucleotide kinase (NEB, M0201L), and 1x T4 ligation buffer under the following conditions: 37°C for 30 minutes, 95°C for 5 minutes, then ramp down to 25°C at 5°C/minute. Duplexed sgRNAs were diluted 1:100, then 2 µL of this dilution was used in a ligation reaction with 100 ng pX-sgRNA-eGFP-MI linearized with BbsI-HF (NEB, R3539S). Each completed sgRNA vector was verified by Sanger sequencing using the human U6 promoter sequencing primer (GGC-CTA-TTT-CCC-ATG-ATT-CC).

### CRISPRi Screen

OVCAR3-KRAB cells were plated at 50,000 cells per well in 24-well plates using antibiotic-free RPMI media supplemented with 10% FBS. After 24 hours, cells were transfected with a total of 300 ng sgRNA vectors using Fugene 6 following the manufacturer’s protocol. Two sgRNAs were designed to target the BRD4 peak summit for each super-enhancer. For negative control wells (empty vector, scramble1, scramble2, Dorm1) and the well targeting the TP53 gene a single sgRNA vector was transfected. For positive control wells (PLAG1 gene promoter, RNF4 gene promoter, FOXL2 gene promoter, RNF4 enhancer, FOXL2 enhancer) and wells targeting each super-enhancer, two sgRNA vectors were co-transfected in each well. Genomic coordinates for all super-enhancers and their sgRNA sequences are found in Supplemental Data 1. 72 hours after transfection of the sgRNAs, cells were visualized for GFP expression to ensure good transfection efficiency. After visualization, wells were washed with 1x PBS and RNA was extracted using the Zymo Quick-RNA Miniprep Kit (Zymo, R1055) with the on-column DNAseI treatment step. RNA-seq libraries were prepared using the Lexogen Quantseq 3’ mRNA-seq FWD Library Prep Kit (Lexogen QuantSeq, 015.2x96) and the PCR Add-On Kit for Illumina (Lexogen QuantSeq, 020.96).

### CRISPRi for SE14 and SE60

OVCAR3-KRAB cells were seeded in 6-well plates at 200,000 cells/well using antibiotic-free RPMI media supplemented with 10% FBS. After 24 hours, cells were transfected with a total of 1.5 µg sgRNA vector per well using Fugene 6 (Promega, E2691) following the manufacturer’s protocol. For negative control wells (Scramble1), a single sgRNA vector was transfected. For wells targeting SE14 and SE60, two unique sgRNAs were co-transfected in each well. 72 hours after transfection, cells were visualized for GFP expression to ensure good transfection efficiency. Cells were then washed with 1x PBS and RNA was extracted using the Zymo Quick-RNA Miniprep Kit (Zymo, R1055) with on-column DNaseI treatment. Experiments were conducted three to four times to ensure reproducibility.

### Super-Enhancer Knockout with CRISPR-Cas9

Targeted deletion of super-enhancers was performed using the CRISPR-cas9 system following published protocols^68–70^. Briefly, guide RNA target sites flanking the BRD4 peak summit for each super-enhancer were selected using the CRISPOR web tool ^47^. Genomic coordinates for all super-enhancers and their sgRNA sequences are found in Supplemental Table 1. Guide RNA oligos were ordered from Integrated DNA Technologies, annealed, and cloned into pSpCas9(BB)-2A-Puro (PX459) V2.0 (Addgene Plasmid #62988). Per super-enhancer targeted, four complete gRNA plasmids (two 5’ and two 3’ of the target site) were transfected into OVCAR3 cells using the Fugene 6 transfection reagent (Promega, E2691) following the manufacturer’s protocol. CRISPR-cas9 positive clones were identified through puromycin selection that began 3 days post-transfection and lasted a total of 7 days. To confirm the deletion of super-enhancer targets, individual positive clones were picked into separate wells and genotyped via PCR using primers flanking the deletion site, along with internal primers used to identify wild type alleles (Supplemental Table 2). Successful SE14 deletion resulted in a ∼2500-2800bp deletion. Successful SE60 deletion resulted in a ∼1700-1800bp deletion. Correct super-enhancer knockout cell lines were further analyzed by Sanger DNA sequencing to determine the precise boundaries of the deletion.

### RNA-seq

For the CRISPRi screen, RNA-seq libraries were prepared using the Lexogen Quantseq 3’ mRNA-seq FWD Library Prep Kit (Lexogen QuantSeq, 015.2x96) and the PCR Add-On Kit for Illumina (Lexogen QuantSeq, 020.96). Libraries underwent 75bp single end sequencing on an Illumina NextSeq 500 instrument (at TGL).

For RNA-seq of OVCAR3 WT, SE60KO1, SE60KO2, SE14KO1rep1, and SE14KO1rep2, libraries were prepared with the Illumina TruSeq Stranded mRNA Kit following the manufacturer’s protocol. Libraries underwent 75bp paired end sequencing on an Illumina NextSeq 500 instrument (at TGL).

For SE60KO3, SE14KO2, SE14KO3, scramble1-KRAB, SE60-KRAB, libraries were created and sequenced by Novogene. These libraries underwent 150bp paired end sequencing on an Illumina NovaSeq 6000 instrument.

### ChIP-seq

OVCAR3-KRAB cells were transfected with sgRNAs targeting either scramble1 (non-targeting) or SE60 (2 pooled sgRNAs) following the same protocol mentioned above for “CRISPRi for SE14 and SE60.” For each of the three replicates conducted, 1-2 million cells were used for fixation with 11% formaldehyde following Active Motif’s Epigenetic Services ChIP Fixation Protocol. ChIP-seq for H3K9me3 was performed by Active Motif using antibody antibody 39161 with spike-in Drosophila normalization. ChIP-seq libraries underwent 75bp single end sequencing on an Illumina NextSeq 5000 instrument by Active Motif.

### Cell Proliferation Assay

Cell collections were performed at Days 0, 2, 4, and 6. Cells were fixed with 10% formaldehyde and stained with a 0.1% crystal violet solution. Incorporated crystal violet was extracted using 10% glacial acetic acid and the absorbance was read at 595 nm. This procedure was conducted four times to ensure reproducibility. Results are shown as the mean OD 595nm reading ± SEM. Statistical analysis was conducted in R using a t-test.

### Cell Migration Assay

OVCAR3 WT and SEKO cells in serum-free RPMI media were seeded to the upper chamber of a transwell insert at 60k cells per insert. The lower chamber contained RPMI with 10% FBS. Cells were incubated for 24 hours, then all non-migrated cells were removed from the upper membrane. Cells were fixed and stained using the Hema 3 Staining Kit (Fisher Scientific, 122-911). Ten brightfield images were taken per insert and images were analyzed using the CellProfiler 4.2.1 software to count the number of cells per transwell-insert. This procedure was conducted four times to ensure reproducibility. Results are shown as the mean cell count ± SEM. Statistical analysis was conducted in R using a t-test.

### General Program Versions

Unless specified these are the versions used for scripting/analysis in R and Python throughout the project for the bulk data analysis of CRISPRi, CRISPR-KO, CNV, and H3K27ac/BRD4 ChIP-Seq data. Unless otherwise stated all “overlap” analysis visualization was performed using intervene ^71^.

**Python:** 3.6.5

**R:** 4.0.0

Intervene: 0.6.5

## RNA Seq: CRISPRi Screen

### General Metrics

RNA-seq was performed following the pipeline put forth by LEXOGEN in the 3’ mRNA-Seq package; namely using STAR, HTSEQ, and DESEQ2. These processes will be explained in more detail below.

### QC

Quality control was performed using the FastQC tool and the results were analyzed (http://www.bioinformatics.babraham.ac.uk/projects/fastqc/). All of the metrics returned as acceptable with no clear failures. We thus proceeded with processing and analysis.

Version: FastQC v0.11.7

### Trimming

Trimming was performed using the bbmap function bbduk.sh with the following parameters (https://sourceforge.net/projects/bbmap/).

Parameters: ktrim=r k=13 useshortkmers=t mink=5 qtrim=r trimq=10 minlength=20 ftm=5 Version: Version 38.46

### Alignment

The trimmed and cleaned reads were then aligned to the HG38v12 human genome using STAR version 2.6.0a with the recommended parameter set and the following conditions ^72^. Parameters:

--runMode alignReads

--genomeDir

--outFilterType BySJout

--outFilterMultimapNmax 20

--alignSJoverhangMin 8

--alignSJDBoverhangMin 1

--outFilterMismatchNmax 999

--outFilterMismatchNoverLmax 0.6

--alignIntronMin 20

--alignIntronMax 1000000

--alignMatesGapMax 1000000

--readFilesCommand gunzip -c

--outSAMtype BAM SortedByCoordinate

--outSAMattributes NH HI NM MD Version: 2.6.0a

### File Formatting

The bam files from STAR were then indexed and sorted using functions in the SAMTOOLS package, namely samtools sort and samtools index ^73^.

Version: 1.9

### Quantification

The sorted and indexed bam files were quantified using htseq and the gencode v29 primary assembly as a reference and with the following parameters ^74^.

Parameters:

-m intersection-nonempty

-s yes

-f bam

-r pos Version: 0.11.2

### Read Distributions

The package RSeQC was used to assess the distribution of reads across the genome. Specifically, the python program read_distribution.py was used with default parameterizations to create a summary of this information ^75^.

Version: 3.0.0

### Review QC

All of the alignment, counting, and cleaning program outputs were assessed with MultiQC for potential issues, of which none were determined ^76^. Default parameters were used. Version: 1.9

### Normalization

The count data was first normalized by removing all of the low count genes (genes with < 1 count in every samples); this data was then read into DESEQ2 ^49^. Within DESEQ2 normalized by scaling and size factors followed by a VST transformation. Batch affects were addressed by utilizing the SVT program (part of the DESEQ2 package) and variation from two surrogate variables was removed for the final analysis.

Version(s): sva_3.38.0, DESeq2_1.30.1 Script: Screen_Preprocessing.R

### Determination of DEGs

Differential gene expression was determined by utilizing a rank-based approach similar to the ranking method used by CMAP for their single replicate screens ^48^. Genes were ranked in order of expression (rank 1 being highest expressed, n being the lowest) within every sample, then all samples were aggregated and a global rank was assigned for every gene.

Next, the change in rank was determined between the within-sample rank and the global rank for every gene in every sample. These changes in rank were used to build a distribution of all rank changes for eFDR analysis.

Script: OVCAR3_Screen_Analysis_with_Plotting_LFC_Comparison.ipynb

### Empirical False Discovery Rate Analysis

Empirical False Discovery Rate, an empirically derived variation of the False Discovery Rate, was determined by choosing a rank change threshold and assessing the median number of genes across controls beyond that threshold as compared to a given sample ^42^. For example, if there are a median of 4 genes in the controls and 40 genes in Sample A; the eFDR for this comparison would be 4/40 or 10%.

Script: OVCAR3_Screen_Analysis_with_Plotting_LFC_Comparison.ipynb

### Relative Expression Correlation Analysis

A log2 fold change was calculated between all genes in a sample and the median of the controls. All genes determined as significant by the rank-based analysis (across all super-enhancers) were aggregated into one pool of genes. This pool of genes was then used to compare RC to LFC values within each super-enhancer to determine the correlation of these sets of values.

Script: OVCAR3_Screen_Analysis_with_Plotting_LFC_Comparison.ipynb

### Clustering

KMeans clustering analysis was used to cluster the differentially ranked gene list. Three clusters were determined as optimal by analysis of the elbow plot and these clusters were then applied to the data. Unsupervised hierarchical clustering was then used to determine the super-enhancer relationships.

Script: OVCAR3_Screen_Analysis_with_Plotting_LFC_Comparison.ipynb

### Pathway Analysis

Genes detected from the differential expression analysis were analyzed using CancerSEA and the molecular signatures database ^50, 77, 78^. This program performs pathway analysis using cell-type specific information relevant to cancer based on available single cell datasets. All of the genes in a given KMeans cluster were fed into this set of programs as a gene list and results were retrieved.

## RNA Seq – CRISPR KO

### General Metrics

RNA-Seq was performed following a similar pipeline to that used in the screen analysis with parameters adjusted to account for differences in the data (paired-end with greater depth); namely using STAR, HTSEQ, and DESEQ2. This will be expounded in more detail below. ***QC*:** Quality control was performed using the FastQC tool (http://www.bioinformatics.babraham.ac.uk/projects/fastqc/). All of the metrics returned as clear or warnings with no failures.

Version: FastQC v0.11.7

### Trimming

No trimming was needed or performed.

### Alignment

The reads were then aligned to the HG38v12 human genome using STAR version 2.6.0a with the following parameter set ^72^.

--runMode alignReads

--outFilterType BySJout

--outFilterMultimapNmax 20

--alignSJoverhangMin 8

--alignSJDBoverhangMin 1

--outFilterMismatchNmax 999

--outFilterMismatchNoverLmax 0.6

--alignIntronMin 20

--alignIntronMax 1000000

--alignMatesGapMax 1000000

--readFilesCommand gunzip -c

--outSAMtype BAM SortedByCoordinate

--outSAMattributes NH HI NM MD

### File Formatting

The bam files from STAR were then indexed and sorted using functions in the SAMTOOLS package, namely samtools sort and samtools index ^73^.

Version: 1.9

### Quantification

The sorted and indexed bam files were quantified using htseq using the gencode v29 primary assembly as a reference and with the following parameters ^74^.

-m union

-nonunique all

-s reverse

--type=gene

--additional-attr=gene_name

-f bam

-r pos gencode.v29.annotation.gff3 Version: 0.11.2

### Read Distributions

The package RSeQC was used to assess the distribution of reads across the genome. Specifically, the python program read_distribution.py was used with default parameterizations to create a summary of this information ^75^.

Version: 3.0.0

### Review QC

All of the alignment, counting, and cleaning program outputs were assessed with MultiQC for potential issues ^76^. Default parameters were used and all of the reports were good. Version: 1.9

### Normalization (Batch Effect Detection)

The count data was first normalized by removing all of the low count genes (genes with < 1 count in every samples); this data was then read into DESEQ2 ^49^. Within the DESEQ framework, the counts data was adjusted for scaling and size factors followed by a VST transformation. Batch affects were addressed by utilizing the SVT program and variation from one surrogate variable was accounted for in the DESEQ2 model.

Version(s): sva_3.38.0, DESeq2_1.30.1

Script: DESEQ2_2021Reps_RNA_SVA_Plotting.Rmd

### Normalization

The pre-VST data was used for standard in-program normalization by DESEQ2 during the differential expression analysis procedure.

Version(s): sva_3.38.0, DESeq2_1.30.1

Script: DESEQ2_2021Reps_RNA_SVA_Plotting.Rmd

### Determination of DEGs

Differential gene expression was determined by utilizing DESEQ2 and default parameters. Genes called as differentially expressed at an FDR adjusted p-value less than 0.0005 were identified and collected for analysis and figure making.

Script: DESEQ2_2021Reps_RNA_SVA_Plotting.Rmd

### Pathway Analysis

Genes detected from the differential expression analysis were analyzed using CancerSEA and the molecular signatures database ^50, 77, 78^. This program performs pathway analysis using cell-type specific information relevant to cancer based on available single cell datasets. The top 100 most significant downregulated genes from differential expression analysis were fed into this program as a gene list and results relevant to ovarian cancer were retrieved.

Script: DESEQ2_2021Reps_RNA_SVA_Plotting.Rmd

### Survival Analysis

To perform survival analyses we made use of the KM plotter tool ^33^. This tool allows a user to look at the effect that expression of induvial genes or a gene set has on overall survival across a number of cancer patients. We looked at the top 100 genes ordered by adjusted P-value (the top 100 most significant genes) as a set (using the median expression of the whole group); and/or looked at genes individually.

## CRISPRi RNA-Seq Analysis

### General Metrics

RNA-Seq was performed following a similar pipeline to that used in the screen analysis with parameters adjusted to account for differences in the data (paired-end with greater depth); namely using STAR, HTSEQ, and DESEQ2. This will be expounded in more detail below.

### QC

Quality control was performed using the FastQC tool (http://www.bioinformatics.babraham.ac.uk/projects/fastqc/). All of the metrics returned as clear or warnings with no failures.

Version: FastQC v0.11.7

### Trimming

No trimming was needed or performed.

### Alignment

The reads were then aligned to the HG38v12 human genome using STAR version 2.6.0a with the following parameter set ^72^.

--runMode alignReads

--outFilterType BySJout

--outFilterMultimapNmax 20

--alignSJoverhangMin 8

--alignSJDBoverhangMin 1

--outFilterMismatchNmax 999

--outFilterMismatchNoverLmax 0.6

--alignIntronMin 20

--alignIntronMax 1000000

--alignMatesGapMax 1000000

--readFilesCommand gunzip -c

--outSAMtype BAM SortedByCoordinate

--outSAMattributes NH HI NM MD

### File Formatting

The bam files from STAR were then indexed and sorted using functions in the SAMTOOLS package, namely samtools sort and samtools index ^73^.

Version: 1.9

### Quantification

The sorted and indexed bam files were quantified using htseq using the gencode v29 primary assembly as a reference and with the following parameters ^74^.

-m union

-nonunique all

-s reverse

--type=gene

--additional-attr=gene_name

-f bam

-r pos gencode.v29.annotation.gff3 Version: 0.11.2

### Read Distributions

The package RSeQC was used to assess the distribution of reads across the genome. Specifically, the python program read_distribution.py was used with default parameterizations to create a summary of this information ^75^.

Version: 3.0.0

### Review QC

All of the alignment, counting, and cleaning program outputs were assessed with MultiQC for potential issues ^76^. Default parameters were used and all of the reports were good. Version: 1.9

### Normalization

The pre-VST data was used for standard in-program normalization by DESEQ2 during the differential expression analysis procedure.

Version(s): sva_3.38.0, DESeq2_1.30.1

Script: DESEQ2_RNA_Plotting_CRISPRi_Analysis_Revised.Rmd

### Determination of DEGs

Differential gene expression was determined by utilizing DESEQ2 and default parameters. Genes called as differentially expressed at a an FDR adjusted p-value less than 0.0005 were identified and collected for analysis and figure making.

Script: DESEQ2_RNA_Plotting_CRISPRi_Analysis_Revised.Rmd

### Pathway Analysis

Genes detected from the differential expression analysis were analyzed using CancerSEA and the molecular signatures database ^50, 77, 78^.

### Survival Analysis

To perform survival analyses we made use of the KM plotter tool ^33^. This tool allows a user to look at the effect that expression of induvial genes or a gene set has on overall survival across a number of cancer patients. We looked at the top 100 genes ordered by adjusted P-value (the top 100 most significant genes) as a set (using the median expression of the whole group); and/or looked at genes individually.

## Copy Number Analysis

### Gathering

The copy number and RNA-seq data for this analysis was downloaded from the TCGA repository Firebrowse (http://firebrowse.org/) which contains the data used in the TCGA analysis of ovarian cancer^11^. We used the TCGA patient barcodes to determine if a tumor was from normal tissue or cancer patients. Samples were subset based on these barcodes to select for tumors. Additionally, for the CNVeQTL analysis, samples unique to each dataset (RNA or Copy Number) were removed. To perform this, we looked for matching patient identifiers between RNA-seq and copy number data and kept any data with ID overlaps.

### Windowing

The autosomal (Chr 1-22) genome (hg19) was divided into 15kb bins using python. We decided to use a sliding window size of 15kb based on the overall size distribution of our super-enhancers. Since the median size of our super-enhancers is 21kb, we wanted a window size similar to the median size but smaller, as smaller windows allow for better resolution. We settled on 15kb as being close to the median size and small enough to give us good resolution, yet large enough to be computationally feasible (smaller window sizes create larger datasets and increase the computational burden of assigning signal and analyzing the data).

Script: Split_Genome_into_windows.ipynb

### Super Enhancer Overlap

Bedtools intersect (one bp overlap) was used to create a subset of the whole genome 15kb sliding windows which overlapped the super-enhancer regions. This gave us two data sets, one being whole genome sliding windows and the other being SE overlapping sliding windows (a subset of the whole genome group).

### Copy Number Assignment - Whole Genome

Patient copy number was assigned to each 15kb window for every chromosome individually using the script OVLP_CNV_Whole_Genome.py. If the patient data overlapped a sliding window by at least one base pair signal from the patient was assigned to this window. Once this was performed for every chromosome individually, the chromosome data was aggregated using Combine_CNV_Chr_Files.ipynb.

Script: OVLP_CNV_Whole_Genome.py & Combine_CNV_Chr_Files.ipynb

### Copy Number Assignment – Super-Enhancer Overlap

Patient copy number was assigned to each 15kb bin for every chromosome individually using the script SEOVLP_CNV.py. If the patient data overlapped a sliding window by at least one base pair signal from the patient was assigned to this window. Chromosome data was aggregated using Combine_CNV_Chr_Files.ipynb.

Script: SEOVLP_CNV.py & Combine_CNV_Chr_Files.ipynb

### CNVeQTL Analysis

Copy Number Expression QTL were identified using MatrixQTL where the SE overlapping CNV windows were defined as the “SNPs” and the matching RNA-Seq data served as the Expression dataset ^41^. Of note, genes with over 100 NA, missing, or 0 values were removed from this dataset prior to analysis. The CNV and RNA data were also converted into float values for ease of use in MatrixEQTL. CNVeQTL were identified using the linear MatrixEQTL algorithm on the original data with a P-value threshold of 1e-3. In order to determine significance, the null hypothesis was induced and used to determine an empirical FDR. The null hypothesis, in which there is no association between specific copy number regions and gene expression, was induced by randomly permuting the column assignments of the RNA-seq data, the CNV data was left alone. This maintains the variance structure of the CNV data and merely changes which CNV data column gets matched with a given RNA-Data column. For example, CNV columns 1,2,3, and 4 (corresponding to patients 1,2,3, and 4) might now be matched with RNA columns 30,75,6, and 210; this allows us to use the same overall data and investigate what happens where there is no link between CNV and RNA values (as patient 1’s CNV values should be random in relation to the gene expression of patient 30). MatrixEQTL was then run on using the original copy number data and the new column shuffled RNA-seq data; this shuffling and running of MatrixEQTL was performed 100,000 times. The median number of significant eQTLs detected across all 100k null conditions was used as the numerator for the empirical false discovery rate analysis, with the experimental results being the denominator. There is some variability in eFDR, as no seed was set and the permutations are random, but all repeats of 100k (3 repeats or 300k trials) returned an eFDR less than 0.1 or 10%.

Script: CNV_eQTL.R

### Determining Super Enhancer Amplification

In order to assess whether the super-enhancer regions were amplified, we compared the distribution of CNV values in the super-enhancer overlapping sliding windows with the whole genome by sub-setting and direct comparison. We performed 10k random subset comparisons, and one direct comparison. In any given comparison, we took the 336 super-enhancer overlapping windows and then randomly drew 336 windows from the whole genome background; these two sets were then compared for significant differences using a Welch’s one-sided t test. This analysis allowed us to determine if the super-enhancer overlapping group was significantly amplified relative to the randomly drawn subset. For the direct comparison, we took all 336 SE overlapping windows and directly compared the CNV values across these windows to the ∼192,000 total regions using the same t test metric.

Script: OVCAR_CNV_Comparison_Final.R

### Survival Analysis

The effect of amplification of these regions on overall survival in patients was calculated using the Kaplan-Meier Log Rank Change test and the Cox Proportional Hazards Model ^32^. The survival data was downloaded from the TCGA and the patient ID was mapped back to the CNV values for each patient ^39^. These datasets were then combined into a single set formatted as described in CNV_KM_Plots.R. This combined survival and copy number dataset was then analyzed using the functions built in CNV_KM_Plots.R to provide a metric of significance for each 15kb copy number region.

Script: CNV_KM_Plots.R

## ChIP Seq (OVCAR3 BRD4 and H3K27ac)

### Data Acquisition

Publicly available ChIP-Seq data was downloaded from the SRA database associated with GSE101408 (experimental OVCAR3 H3K27ac condition) using fastq dump ^29^. This process was repeated to get BRD4 binding data for DMSO treated OVCAR3 cells as well as the input control from GSE77568 ^30^.

### Processing

The following steps were used to process each file separately (H3K27ac, BRD4 ChIP, and BRD4 sample input). At the peak calling step, the BRD4 ChIP data was informed by the processed input control. As there was no input provided for the H3K27ac data, no input was processed or utilized for this sample.

### Data Quality Check

The quality of the data was assessed using fastqc and reads were trimmed using Trimmomatic (version 0.38) with the following parameters ^79^.

Leading: 30

Trailing: 30

Sliding Window: 4:30

MINLEN: 36

Phred33

### Alignment

The fastq files were aligned to hg19 using Bowtie2 with default parameters ^80^. The output sam files were then converted to bam files using samtools and sorted/indexed.

### Processing Bam Files (Marking Duplicates)

The aligned and sorted bam files were then marked for duplicate reads using picard with the following parameters ^81^.

java -Xmx4G -jar $PICARD/picard.jar MarkDuplicates

VALIDATION_STRINGENCY=LENIENT

ASSUME_SORTED=true

REMOVE_DUPLICATES=false

### Processing Bam Files (Removing Duplicates)

The duplicate reads marked by Picard were then removed by samtools using the following command.

samtools view -F 1804 -b in.bam > clean.bam

### Create tagAlign Files

A tagAlign file was generated using the following command.

bamToBed -i clean.bam | awk ‘BEGIN{OFS=“\t”}{$4=“N”;$5=“1000”;print $0}’ | tee clean.tagAlign | gzip -c > clean.tagAlign.gz

### Peak Calling

Peaks were identified using MACS2 with the following parameterization ^34^. The input sample was used as the control for the BRD4 ChIP data; the H3K27ac data was processed without an input with MACS2 determining the control by default processes.

Version: 2.2.6

*BRD4*:

-g hs

-p 1e-2

--nomodel

--extsize 121

-B

*H3K27ac*:

-g hs

-p 1e-2

--nomodel

--extsize 218

-B

### Determination of the Final Peak Set

The called peaks were then intersected with all genes in the hg19 human genome, using bedtools intersect (1bp overlap) and overlapping regions were removed (https://bedtools.readthedocs.io/en/latest/). The remaining peaks from H3K27ac and BRD4 that did not overlap genes were then intersected using bedtools (1bp overlap) and regions with both an H3K27ac and BRD4 peak were kept (using the BRD4 coordinates).

### Creating BigWigs

The fold enrichment of the bam files were calculated across these peaks for both H3K27ac and BRD4 using macs2 bdgcmp and the -m ppois parameter. As the H3K27ac had no input we felt that in order to allow for fair comparison both H3K27ac and BRD4 bedgraph files should use the -m ppois parameter (we did also generate a fold enrichment aka FE bedgraph for BRD4 to ensure it was comparable to the ppois version). These bedgraph files were then converted to bigwigs using bedGraphToBigWig from UCSC.

### Calling Super Enhancers

Super Enhancers were then identified using the ROSE ^36^ pipeline with default parameters.

Version: 0.1

Python: 2.7

### Meta-Analysis

Meta plots and heatmaps for these data were created using Deeptools. We generated matrices using signal from the bigwig files and the overlapping 12,339 peaks as the regions. These matrices were then used for plotting.

Version: 3.1.0

### ChIP-Seq (H3K9me3)

ChIP-Seq analysis for H3K9me3 was performed by ACTIVEMOTIF following their spike in protocol, the following is a modified excerpt from the workflow provided to us.

### Sequence Analysis

The 75-nt single-end (SE75) sequence reads generated by Illumina sequencing (using NextSeq 500) were mapped to the genome using the BWA algorithm (“bwa aln/samse” with default settings). Alignment information for each read is stored in the BAM format. Only reads that pass Illumina’s purity filter, align with no more than 2 mismatches, and map uniquely to the genome were used in the subsequent analysis. In addition, duplicate reads (“PCR duplicates”) were removed.

### Determination of Fragment Density

Since the 5′-ends of the aligned reads (= “tags”) represent the end of ChIP/IP-fragments, the tags were extended in silico (using Active Motif software) at their 3′-ends to a length of 200 bp, which corresponds to the average fragment length in the size-selected library. To identify the density of fragments (extended tags) along the genome, the genome was divided into 32-nt bins and the number of fragments in each bin is determined. This information (“signal map”; histogram of fragment densities) is stored in a bigWig file. bigWig files also provide the peak metrics in the Active Motif analysis program described below.

### Peak Finding

The generic term “Interval” is used to describe genomic regions with local enrichments in tag numbers. Intervals are defined by the chromosome number and a start and end coordinate. The peak caller used at Active Motif for this project was SICER^82^. This method was used to detect significant enrichments in the ChIP/IP data file when compared to the Input data file or relative to neighboring background regions.

### Additional Analysis Steps

***a. Standard Normalization***: In the default analysis, the tag number of all samples (within a comparison group) is reduced by random sampling to the number of tags present in the smallest sample.

***b. Spike-in Adjustment***: Spike-in of Drosophila chromatin was performed; the number of test tags were adjusted (again by down-sampling) by a factor that would result in the same number of spike-in Drosophila tags for each sample.

### Merged Region Analysis

To compare peak metrics between 2 or more samples, overlapping Intervals (orange bars in diagram below) were grouped into “Merged Regions” (green bars), which are defined by the start coordinate of the most upstream Interval and the end coordinate of the most downstream Interval (= union of overlapping Intervals; “merged peaks”). In locations where only one sample has an Interval, this Interval defines the Merged Region. The use of Merged Regions was necessary because the locations and lengths of Intervals are rarely exactly the same when comparing different samples. Furthermore, with this approach fragment density values could be obtained even for samples for which no peak was called.

### Annotations

After defining the Intervals and Merged Regions, their genomic locations along with their proximities to gene annotations and other genomic features are determined. In addition, average and peak (i.e. at “summit”) fragment densities within Intervals and Merged Regions were compiled.

### Differential Binding Analysis

DESeq2 was used to determine regions of differential binding.

**Hi-C**

### In Situ Hi-C

OVCAR3 cells were grown under recommended culture conditions in RPMI media supplemented with 10% FBS and 1% penicillin/streptomycin. Four to five million cells were fixed with 1% formaldehyde for 10 minutes. Pellets were flash frozen in liquid nitrogen and stored at -80°C.

In situ Hi-C was performed as previously described ^83^. Pellets were lysed in ice-cold Hi-C lysis buffer (10mM Tris-HCl pH 8.0, 10mM NaCl, 0.2% IGEPAL CA630) with 50μL of protease inhibitors for 15 min on ice. Cells were pelleted and washed once more using the same buffer. Pellets were resuspended in 50μL of 0.5% SDS and incubated at 62°C for 7 min. Reactions were quenched with 145μL water and 25μL 10% Triton X-100 at 37°C for 15 min. Chromatin was digested overnight with 25μL of 10X NEBuffer2 and 100U of MboI at 37°C with rotation.

Reactions were incubated at 62°C for 20 min to inactivate MboI, then cooled to RT. Fragment overhangs were repaired by adding 37.5μL 0.4mM biotin-14-dATP; 1.5μL each 10mM dCTP, dGTP, dTTP; 8μL 5U/μL DNA Polymerase I, Large (Klenow) Fragment and incubating at 37°C for 1.5 h with rotation. Ligation was performed by adding 673μL water, 120μL 10X NEB T4 DNA ligase buffer, 100μL 10% Triton X-100, 6μL 20mg/mL BSA, and 1μL 2000U/μL T4 DNA ligase and incubating at RT for 4 h with slow rotation. Samples were pelleted at 2500*g*, resuspended in 432μL water, 18μL 20mg/mL proteinase K, 50μL 10% SDS, and 46μL 5M NaCl, incubated at 55°C for 30 min, and then transferred to 68°C overnight.

Samples were cooled to RT and 1.6x volumes of pure ethanol and 0.1x volumes of 3M sodium acetate pH 5.2 were added to each sample, which were subsequently incubated at -80°C for over 4-6 h. Samples were spun at max speed at 2°C for 15 min and washed twice with 70% ethanol. The resulting pellet was dissolved in 130μL of 10mM Tris-HCl pH 8.0 and incubated at 37°C for 1-2 h. Samples were stored at 4°C overnight.

DNA was sheared using the Covaris LE220 (Covaris, Woburn, MA) to a fragment size of 300-500bp in a Covaris microTUBE. DNA was transferred to a fresh tube and the Covaris microTUBE was rinsed with 70μL of water and added to the sample. A 1:5 dilution of DNA was run on a 2% agarose gel to verify successful shearing.

Sheared DNA was size selected using AMPure XP beads. 0.55x volumes of 2X concentrated AMPure XP beads were added to each reaction and incubated at RT for 5 min. Beads were reclaimed on a magnet and the supernatant was transferred to a fresh tube. 30μL of 2X concentrated AMPure XP beads were added and incubated for 5 min at RT. Beads were reclaimed on a magnet and washed with fresh 70% ethanol. Beads were dried for 5 min at RT prior to DNA elution in 300μL of 10mM Tris-HCl pH 8. Undiluted DNA was run on a 2% agarose gel to verify successful size selection between 300-500 bp.

150μL of 10mg/mL Dynabeads MyOne Streptavidin T1 beads were washed with 400μL of 1X Tween washing buffer (TWB; 250μL Tris-HCl pH 7.5, 50μL 0.5M EDTA, 10mL 5M NaCl, 25μL Tween 20, 39.675μL water). Beads were then resuspended in 300μL of 2X Binding Buffer (500μL Tris-HCl (pH 7.5), 100μL 0.5M EDTA, 20mL 5M NaCl, 29.4mL water), added to the DNA sample, and incubated at RT for 15 min with rotation. DNA-bound beads were then washed twice with 600μL of 1X TWB at 55°C for 2 min with shaking. Beads were resuspended in 100μL 1X NEBuffer T4 DNA ligase buffer, transferred to a new tube, and reclaimed.

Sheared ends were repaired by resuspending the beads in 88μL of 1X NEB T4 DNA Ligase Buffer with 1mM ATP, 2μL of 25mM dNTP mix, 5μL of 10U/uL NEB T4 PNK, 4uL of 3U/uL NEB T4 DNA polymerase I, and 1uL of 5U/uL NEB DNA polymerase 1, large (Klenow) fragment and incubating at RT for 30 min. Beads were washed two more times with 1X TWB for 2 min at 55°C with shaking. Beads were washed once with 100uL of 1X NEBuffer 2, transferred to a new tube, and resuspended in 90uL of 1X NEBuffer 2, 5uL of 10mM dATP, and 5uL of NEB Klenow exo minus, and incubated at 37°C for 30 min. Beads were washed two more times with 1X TWB for 2 min at 55°C with shaking. Beads were washed in 100uL 1X Quick Ligation Reaction Buffer, transferred to a new tube, reclaimed, and resuspended in 50uL of 1X NEB Quick Ligation Reaction Buffer. 2uL of NEB DNA Quick Ligase and 3uL of an appropriate Illumina indexed adapter (TruSeq nano) were added to each sample before incubating at RT for 15 minutes. Beads were reclaimed and washed twice with 1X TWB for 2 min at 55°C. Beads were washed in 100uL 10mM Tris-HCl pH 8, transferred to a new tube, reclaimed, and resuspended in 50uL of 10mM Tris-HCl pH 8.

Hi-C libraries were amplified directly off T1 beads with 10 cycles in 5uL of PCR primer cocktail, 20uL of Enhanced PCR mix, and 25uL of DNA on beads. The PCR settings were as follows: 3 min at 95°C followed by 4-12 cycles of 20s 98°C, 15s at 60°C, and 30s at 72°C. Samples were held at 72°C for 5 min before lowering for holding at 4°C. Amplified samples were transferred to a new tube and brought to 250uL in 10mM Tris-HCl pH 8.

Beads were reclaimed and the supernatant containing the amplified library was transferred to a new tube. Beads were resuspended in 25uL of 10mM Tris-HCl pH 8 and stored at -20°C. 0.7x volumes of warmed AMPure XP beads were added to the supernatant sample and incubated at RT for 5 min. Beads were reclaimed and washed once with 70% ethanol without mixing. Ethanol was aspirated. Beads were resuspended in 100uL of 10mM Tris-HCl pH 8, 70uL of fresh AMPure XP beads were added, and the solution was incubated for 5 min at RT. Beads were reclaimed and washed twice with 70% ethanol without mixing. Beads were left to dry and DNA was eluted in 25uL of 10mM Tris-HCl pH 8. The resulting libraries were next quantified by Qubit and Tapestation. A low depth sequence was performed first using the Miniseq sequencer system (Illumina) and analyzed using the Juicer pipeline to assess quality. The resulting libraries underwent paired-end 2x150bp sequencing on an Illumina NovaSeq sequencer. Each replicate was sequenced to an approximate depth of 730 million reads. The full sequencing depth was approximately 2.92 billion reads.

### Hi-C Data Processing and Analysis

In situ Hi-C datasets were processed using a modified version of the Juicer Hi-C pipeline (https://github.com/EricSDavis/dietJuicer) with default parameters as previously described ^84^. MboI was used as the restriction enzyme, and reads were aligned to the hg19 human reference genome with bwa (version 0.7.17). Four biological replicates were aligned and merged for a total of 2,922,558,308 Hi-C read pairs in OVCAR3 cells yielding 2,598,024,810 valid Hi-C contacts (88.90%). For visualization, the resulting Hi-C contact matrix was normalized with the “KR” matrix balancing algorithm as previously described to adjust for regional background differences in chromatin accessibility^85^.

Hi-C contact frequency was used to classify CRISPR-KO gene targets as direct or indirect. We compared the fold-change in observed over expected contact frequency between SE14 or SE60 and their respective gene targets with 100 permutations of distance-matched region-gene pairs as controls. Direct targets were defined as SE-gene pairs with an observed/expected contact frequency greater than the 75th percentile of the control distribution. Since distance-matching is only relevant for regions within a chromosome, we restricted our analysis to intra-chromosomal pairs. We performed this analysis on 1) CRISPR-KO-validated target genes and 2) significantly down-regulated (LFC < -0.5) CRISPR-KO-validated target genes. The analysis was conducted in R (4.1.0) using the following R/Bioconductor packages: *GenomicRanges* (1.45.0), *data.table* (1.14.2), *Homo.sapiens* (1.3.1), *InteractionSet* (1.21.1), *plyranges* (1.13.1), *ggplot2* (3.3.5), *ggrepel* (0.9.1)^86^. Example regions were visualized with the *plotgardener* (1.0.3) Bioconductor package. Scripts can be made available upon request^87^.

## Single-cell analysis

### Data Acquisition

We obtained the single-cell RNA-seq and single-cell ATAC-seq data from the GEO accession number GSE173682.

### scRNA-seq Data Processing and Barcode Quality-Control (QC)

For each patient tumor sample, the filtered feature-barcode matrix was converted into a Seurat object using the Seurat R package (Seurat version 3.2) ^88^. To enrich for high quality cells in each patient dataset, QC and doublet removal were performed for each patient dataset individually. First, outlier cells were defined in each of the following metrics: log(UMI counts) (>2 MADs, low end), log(number of genes expressed) (>2 MADs, low end) and log(percent mitochondrial read count +1) (>2 MADs, high end). Only cells meeting all three criteria were kept for doublet detection. To reduce the false positive rate in doublet calling, only cells marked as doublets by both DoubletDecon ^89^(version 1.1.5) and DoubletFinder ^90^ (version 2.0.3) were removed from downstream analysis. After QC and doublet removal for each patient dataset, the individual patient datasets were combined using Seurat’s merge().

### scRNA-seq Clustering and Cell Type Annotation

The merged gene expression matrix was normalized using Seurat’s NormalizeData() with the normalization method set to “LogNormalize.” Feature selection was performed with Seurat’s FindVariableFeatures() with the selection method set to “vst” and the number of top variable features set to 2,000. Prior to principal component analysis (PCA), we scaled the expression for all genes in the dataset using Seurat’s ScaleData(). The top 2,000 most variable genes were summarized by PCA into 50 principal components (PCs) and the cells were visualized in a two-dimensional UMAP embedding using Seurat’s RunUMAP() with all 50 PCs, as suggested by the results of Seurat’s JackStraw() (data not shown). To identify groups of transcriptionally distinct cells, graph-based Louvain clustering was performed using Seurat’s FindNeighbors() with all 50 PCs and Seurat’s FindClusters() with a resolution of 0.7. scRNA-seq UMAP plots were generated in R using ggplot2.

Cell type annotation was performed using a combination of 1) reference-based annotation with the R package SingleR ^91^ and 2) gene signature enrichment with Seurat’s *AddModuleScore()*. After QC, doublet removal, and dimension reduction for each patient dataset, single cells were annotated to known cell types using SingleR with a reference scRNA-seq dataset. Both scRNA-seq datasets were annotated based on a reference scRNA-seq dataset from a human ovarian tumor (sample ID: HTAP*P-*624-SM*P-*3212) ^92^. The individual patient datasets were then combined using Seurat’s *merge()* to form each patient cohort presented in this study and subsequently reprocessed according to the normalization, feature selection and clustering methods described previously. The resulting clusters in each patient cohort dataset were annotated based on the majority cell type label within each cluster. Finally, SingleR cell type annotations were verified by calculating single cell enrichment scores for cell type gene signatures from PanglaoDB ^93^ using Seurat’s *AddModuleScore()*. The cell type annotations for each cluster were then modified to include the cluster number identity hyphened with the cell type identity. These defined the final cell type subcluster identities for scRNA-seq that were used in label transferring to the matching scATAC-seq data.

### scRNA-seq Differential Gene Expression Analysis

Differential gene expression was computed using Seurat’s *FindMarkers()* with the “test.use” parameter set to “wilcox” for the Wilcoxon Rank Sum test. Genes with a Bonferroni-corrected p-value <= 0.01 & average logFC >= 0.1 were deemed upregulated in the cancer epithelial fraction relative to the remaining cell type clusters.

### scATAC-seq Data Processing and Barcode Quality-Control (QC)

The scATAC-seq fragments file for each patient tumor sample was read into the R package ArchR (version 0.9.3) to perform quality control and doublet removal for each patient dataset individually ^94^. To enrich for cellular barcodes, we took advantage of the bimodal distributions in log10(TSS enrichement+1) and in log10(number of unique fragments) characterizing two different populations of barcodes (cellular and non-cellular). Barcode cutoff thresholds for log10(TSS enrichement+1) and log10(number of unique fragments) were estimated using a Gaussian Mixture Model (GMM) for each metric, as implemented in the R package mclust ^95^. Only barcodes above these estimated thresholds in both metrics were kept as cellular barcodes for doublet detection. Doublet enrichment scores were calculated for cellular barcodes using ArchR’s addDoubletScores() with the knnMethod set to “UMAP.” Cellular barcodes with doublet enrichment scores >1 were deemed as putative doublets and subsequently removed using ArchR’s filterDoublets().

### scRNA-seq Cell Type Label Transfer to scATAC-seq

Before transferring labels from scRNA-seq to scATAC-seq, gene activity scores were inferred in scATAC-seq using ArchR’s *addGeneScoreMatrix()*. Briefly, this method uses the following features to estimate gene activity: 1) fragment counts mapping to the gene body, 2) an exponential weighting function to give higher weights to fragment counts closer to the gene and lower weights to fragment counts father away from the gene, and 3) gene boundaries to prevent the contribution of fragments from other genes. Seurat’s CCA implementation in *FindTransferAnchors()* and *TransferData()* was used to assign each of the scATAC-seq cells a cell type subcluster identity from the matching scRNA-seq data and an associated label prediction score ^88^. This label transferring procedure was constrained to only align cells of the same patient dataset (e.g. Patient 1 scATAC-seq cells were assigned only to cell type subclusters represented by Patient 1 scRNA-seq cells). All scATAC-seq cells were included in UMAP visualization, but only scATAC-seq cells with a label prediction score >0.5 were included in downstream analyses. Also, only inferred cell type subclusters with >30 cells were included in downstream analysis to ensure enough cells for downstream analysis.

### scATAC-seq Peak Calling and Data Visualization

After scATAC-seq cells received a cell type subcluster label, pseudo-bulk replicates were generated for each inferred cell type subcluster in the R package ArchR and pseudo-bulk peak calling was performed within each inferred cell type subcluster using MACS2 ^34, 94^. ArchR’s default iterative overlap procedure was used to merge all peak calls into a single peak by barcode matrix across all cellular barcodes in the merged scATAC-seq dataset. Browser tracks visualizing the scATAC-seq coverage per inferred cell type were generated using ArchR’s plotBrowserTrack function.

### scATAC-seq Differential Peak Accessibility for Determining Cancer-Enriched Enhancers

Differential peak accessibility was computed with ArchR’s *getMarkerFeatures()* with the bias argument set to include both “TSSEnrichment” and “log10(number of fragments).” This procedure identifies differentially accessibility peaks (DEPs) between two groups of cells using a Wilcoxon Rank Sum test. DEPs were identified for each cell cluster by comparing the accessibility values of peaks across all cells in a cluster (group 1) relative to the accessibility values for a group of background cells matched for TSS enrichment and read depth (group 2). Peaks with Benjamini-Hochberg FDR <= 0.10 & Log2FC >= 0.25 were deemed cancer-enriched with statistically significant increased accessibility in the cancer epithelial fraction relative to the remaining cell type clusters.

### Enhancer Motif Analysis in scATAC-seq

The sequences of the select cancer-enriched enhancers were extracted with Bedtools getfasta() using the hg38 reference genome ^96^. The enhancer sequences were inputted into FIMO motif scanning with default parameters using a motif database supplied by JASPAR2020 ^56^ ^97^. The FIMO output listed matching motif occurrences with p-value <1e-4. This list was further sorted by Benjamini-Hochberg corrected q-values and TF expression in the cancer fraction by summing the normalized TF counts across all cells within the cancer epithelial clusters. TF expression violin plots were generated with Seurat’s VlnPlot() function.

## DATA AVAILABILITY

Data generated in this study are publicly available in the Gene Expression Omnibus (https://www.ncbi.nlm.nih.gov/geo/) under the accession number GSE174259 (reviewer token sjelgugwrzghlor). The single cell genomics data were downloaded from GEO accession number GSE173682.

## CODE AVAILABILITY

Programs and Scripts for the bulk data analysis, mentioned in the methods, are located at the Github repository: https://github.com/mkelly9513/OV-Project-One

Programs and scripts for the single cell data analysis are located at the Github repository: https://github.com/RegnerM2015/scOVAR_SE_Screen

Programs and scripts for the Hi-C data analysis are available on request to https://github.com/EricSDavis (no “original” scripts were used).

## Supporting information

List_of_Supplemental_Data

## ACKNOWLEGEMENTS

We thank Dr. Victoria Bae-Jump for her helpful clinical insights on ovarian cancer. We also thank the members of the Franco Lab for their helpful comments and discussions along the way. This work was funded by grants from the NIH/National Cancer Institute (5-P50-CA058223-25), the Susan G. Komen Breast Cancer Research Foundation (CCR19608601), and the V Foundation for Cancer Research (V2019-015) to H.L.F. Additional support was provided by NIH grants 5T32-CA217824 (Cancer Epigenetics Training Program) to A.A.P. and M.W.L., NIGMS Training grant T32-GM067553 to E.S.D., and NIH grant R35-GM128645 to D.H.P.

## AUTHOR CONTRIBUTIONS

H.L.F and M.R.K conceived and supervised the study with input from K.W. The computational analyses were designed and performed by M.R.K. with input from M.J.R., J.S.P., and H.L.F. The CRISPR based experiments and molecular biology was performed by K.W. with help from M.W.L. The Hi-C data was generated and analyzed by A.A.P., E.S.D., and D.H.P. The manuscript was written by M.R.K., K.W., and H.L.F. with input from all authors.

## DECLARATION ON INTERESTS

The authors declare no competing interests

## Notes

### Competing Interest Statement

The authors have declared no competing interest.

### Summary of Updates

Author affiliations updated, orcid's linked

