## Supplementary material for "A multi-omic dissection of super-enhancer driven oncogenic gene expression programs in ovarian cancer": Supplimental_Figures

**This PDF file contains the following Supplemental Information:**

Supplemental Figures 1-9

Supplemental Tables 1-2

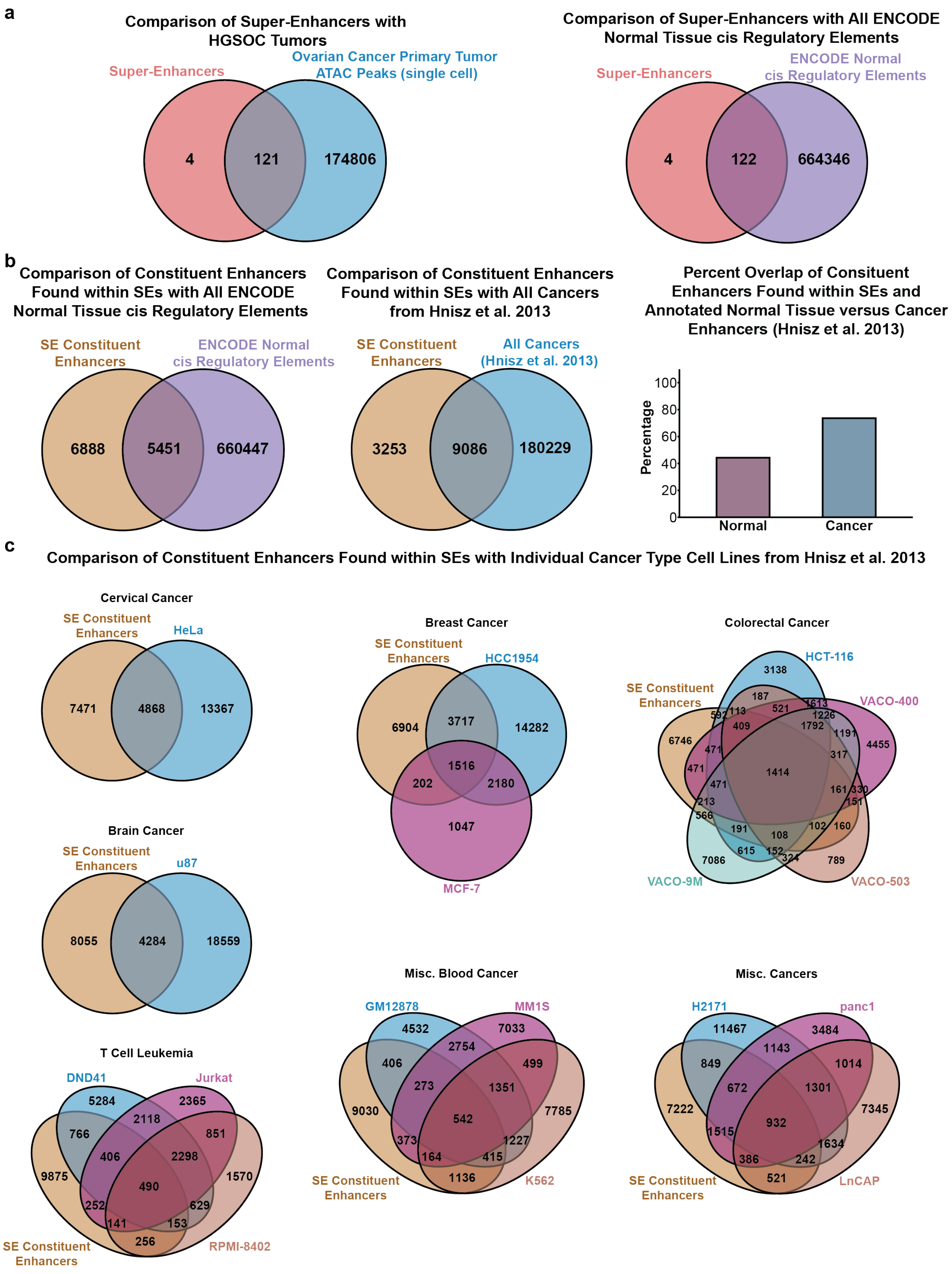
a
Comparison of Super-Enhancers with HGSOC Tumors
Super-Enhancers
Ovarian Cancer Primary Tumor ATAC Peaks (single cell)
4
121
174806
Comparison of Super-Enhancers with All ENCODE Normal Tissue cis Regulatory Elements
Super-Enhancers
ENCODE Normal cis Regulatory Elements
4
122
664346
b
Comparison of Constituent Enhancers Found within SEs with All ENCODE Normal Tissue cis Regulatory Elements
SE Constituent Enhancers
ENCODE Normal cis Regulatory Elements
6888
5451
660447
Comparison of Constituent Enhancers Found within SEs with All Cancers from Hnisz et al. 2013
SE Constituent Enhancers
All Cancers (Hnisz et al. 2013)
3253
9086
180229
Percent Overlap of Consituent Enhancers Found within SEs and Annotated Normal Tissue versus Cancer Enhancers (Hnisz et al. 2013)
Percentage
0
20
40
60
80
100
Normal
Cancer
c
Comparison of Constituent Enhancers Found within SEs with Individual Cancer Type Cell Lines from Hnisz et al. 2013
Cervical Cancer
SE Constituent Enhancers
HeLa
7471
4868
13367
Brain Cancer
SE Constituent Enhancers
u87
8055
4284
18559
T Cell Leukemia
DND41
Jurkat
5284
2365
2118
766
851
406
2298
9875
490
1570
252
629
141
153
256
SE Constituent Enhancers
RPMI-8402
Breast Cancer
SE Constituent Enhancers
HCC1954
6904
3717
14282
1516
202
2180
1047
MCF-7
Colorectal Cancer
HCT-116
SE Constituent Enhancers
VACO-400
3138
187
592
113
521
613
1226
409
1792
1191
6746
4455
471
317
471
1414
471
161
330
213
151
566
191
108
102
160
615
152
324
7086
789
VACO-9M
VACO-503
Misc. Blood Cancer
GM12878
MM1S
4532
7033
2754
406
499
273
1351
9030
542
7785
373
1227
164
415
1136
SE Constituent Enhancers
K562
Misc. Cancers
H2171
panc1
11467
3484
1143
849
1014
672
1301
7222
932
7345
1515
1634
386
242
521
SE Constituent Enhancers
LnCAP

**Supplemental Figure 1. OVCAR3 Super-Enhancers Show Enrichment of Activity in HGSOC Patient Tumors and Cancerous Tissues**

- **a.** Overlap of hg38 detected SEs (lifted-over from Figure 1) with enhancers detected in the cancer cell fraction of HGSOC tumors (*left*). Overlap of hg19 detected SEs (from Figure 1) with all enhancer elements contained in the ENCODE database (*right*).
- **b.** Overlap of constituent enhancers found within SEs (Figure 1) and all normal tissue cis regulatory elements in ENCODE (*left*), and all cancer associated enhancers documented in the Hnisz et al. 2013 research article (*middle*). Breakdown of the percentage of the constituent enhancers found in normal cells (purple) vs cancer cells (blue) (*right*).
- **c.** Overlap of constituent enhancers found within SEs (Figure 1) and all cancer associated enhancers documented in the Hnisz et al. 2013 research article broken down by cancer type.

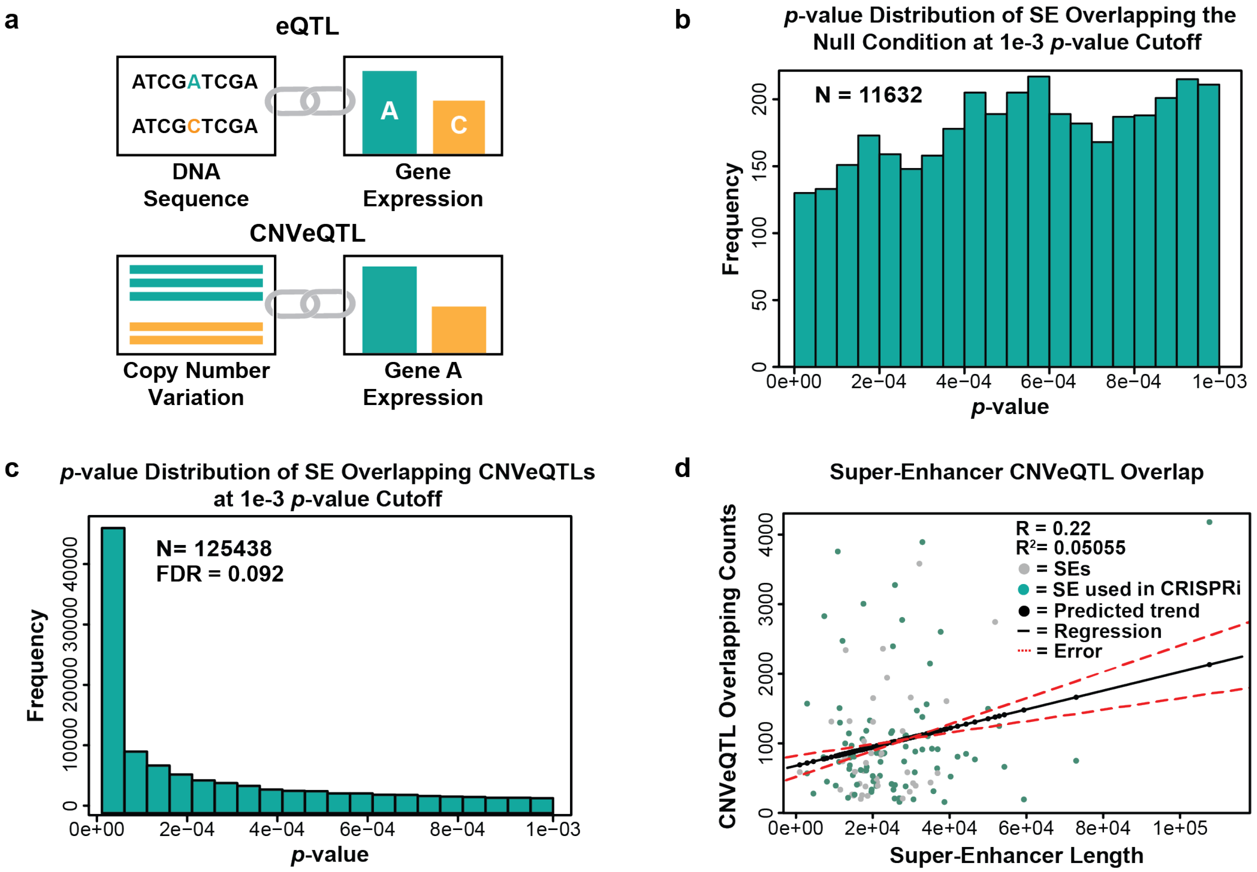


**Supplemental Figure 2: Copy Number Expression Quantitative Loci Analysis (CNVeQTL) in Ovarian Cancer Patients Nominates Salient Super-Enhancers.**

- **a.** Cartoon comparing expression-based quantitative trait loci (eQTL) analysis versus copy-number-based expression quantitative trait loci (CNVeQTL) analysis.
- **b.** Histogram of CNVeQTL *p*-values from one of the 100k permuted null distributions used to calculate the empirically derived false discovery rate (eFDR) of the SE overlapping CNVeQTLs. Median number of CNVeQTLs at *p*-value threshold of $1x10^{-3}$ across all 100k null distributions is shown (n = 11,632).
- **c.** Histogram of CNVeQTL *p*-values from the experimental distribution (SE overlapping CNVeQTLs). The eFDR was determined by comparing the amount of significant CNVeQTL in the experimental group versus the median CNVeQTL count of the 100k permuted nulls.
- **d.** Scatter plot showing the relationship between SE length (in base pairs) and the number of significantly associated CNVeQTLs. The direct association between genomic size and CNVeQTL count shown is by the regression line in black with error bounds depicted as dashed red lines. Dots that represent the SEs that were included into the CRISPRi screen are colored in teal.

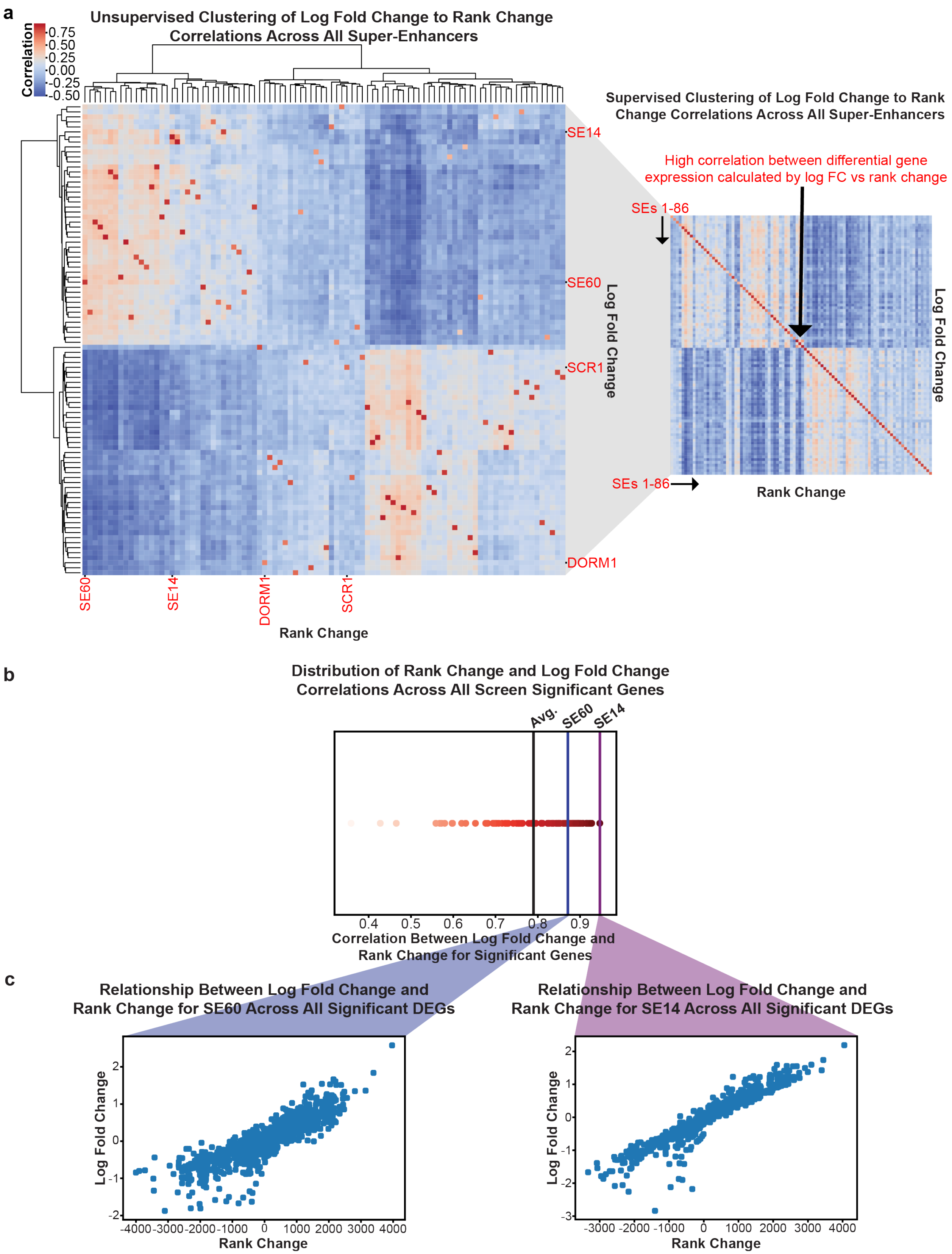


**Supplemental Figure 3: Expression Analysis of the CRISPRi Screen Reveals High Correlation with Rank Change Values**

- **a.** Unsupervised hierarchically clustered heatmap showing the relationship between Log2 Fold Change and Rank Change values for all super-enhancers and their detected target genes. Denoted in red are SE60 and 14 as well as controls (*left*). The same correlation data is provided in a supervised heatmap where the LFC and Rank Change for every SE follow the paired diagonal (*right*).
- **b.** Distribution of all Rank Change to LFC correlations for every SE. The average correlation (between Rank Change and LFC) of all SEs is denoted in black, the correlation between LFC and RC values for SE60 significant DEGs is denoted in blue, and SE14 is denoted in purple.
- **c.** Scatterplot showing the relationship between rank change and log2 fold change for all detected DEGs (from any super-enhancer) genes in SE60 (*left*) and SE14 (*right*).

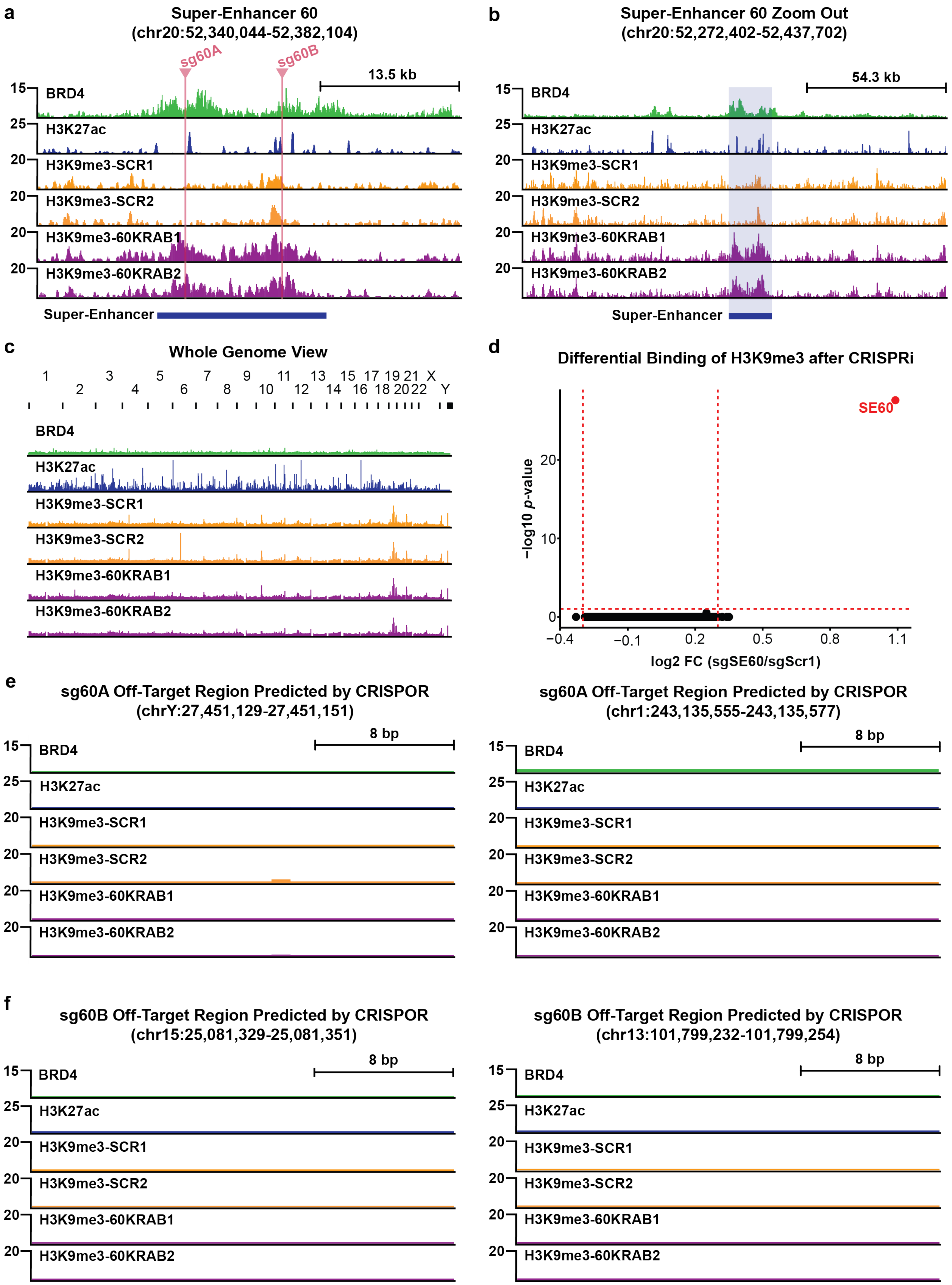
a
Super-Enhancer 60
(chr20:52,340,044-52,382,104)
sg60A
sg60B
13.5 kb
BRD4
H3K27ac
H3K9me3-SCR1
H3K9me3-SCR2
H3K9me3-60KRAB1
H3K9me3-60KRAB2
Super-Enhancer
b
Super-Enhancer 60 Zoom Out
(chr20:52,272,402-52,437,702)
54.3 kb
c
Whole Genome View
d
Differential Binding of H3K9me3 after CRISPRi
SE60
−log10 p-value
log2 FC (sgSE60/sgScr1)
e
sg60A Off-Target Region Predicted by CRISPOR
(chrY:27,451,129-27,451,151)
sg60A Off-Target Region Predicted by CRISPOR
(chr1:243,135,555-243,135,577)
8 bp
f
sg60B Off-Target Region Predicted by CRISPOR
(chr15:25,081,329-25,081,351)
sg60B Off-Target Region Predicted by CRISPOR
(chr13:101,799,232-101,799,254)

**Supplemental Figure 4: H3K9me3 ChIP-seq Reinforces Targeted CRISPRi of Super-Enhancer 60**

- **a.** Browser track view of the SE60 locus investigating the amount of H3K9me3 in scramble negative controls and SE60-KRAB samples. sgRNA target sites are labeled in pink. There is an enrichment in H3K9me3 with the SE60-KRAB condition relative to the scramble controls.
- **b.** Zoomed out browser track view of the SE60 locus investigating the amount of H3K9me3 in scramble controls and SE60-KRAB samples. There is an enrichment in the SE60-KRAB condition relative to the scramble controls that covers ~20kb around the SE60 sgRNA target sites.
- **c.** Browser track of the entire human genome investigating the amount of H3K9me3 in scramble controls and SE60-KRAB samples. There are no clear differences between scramble and SE60-KRAB samples.
- **d.** Volcano plot depicting the results of the differential binding analysis between scramble guide controls and SE60-KRAB targeted knockdown samples, significant regions are denoted in red.
- **e.** Browser track view investigating the amount of H3K9me3 in scramble controls and SE60-KRAB sample for the first (*left*) and second (*right*) highest predicted off-targets of the SE60A sgRNA. There are no clear differences between scramble and SE60-KRAB samples.
- **f.** Browser track view investigating the amount of H3K9me3 in scramble controls and SE60-KRAB samples for the first (*left*) and second (*right*) highest predicted off-targets of the SE60B guide. There are no clear differences between scramble and SE60-KRAB samples.

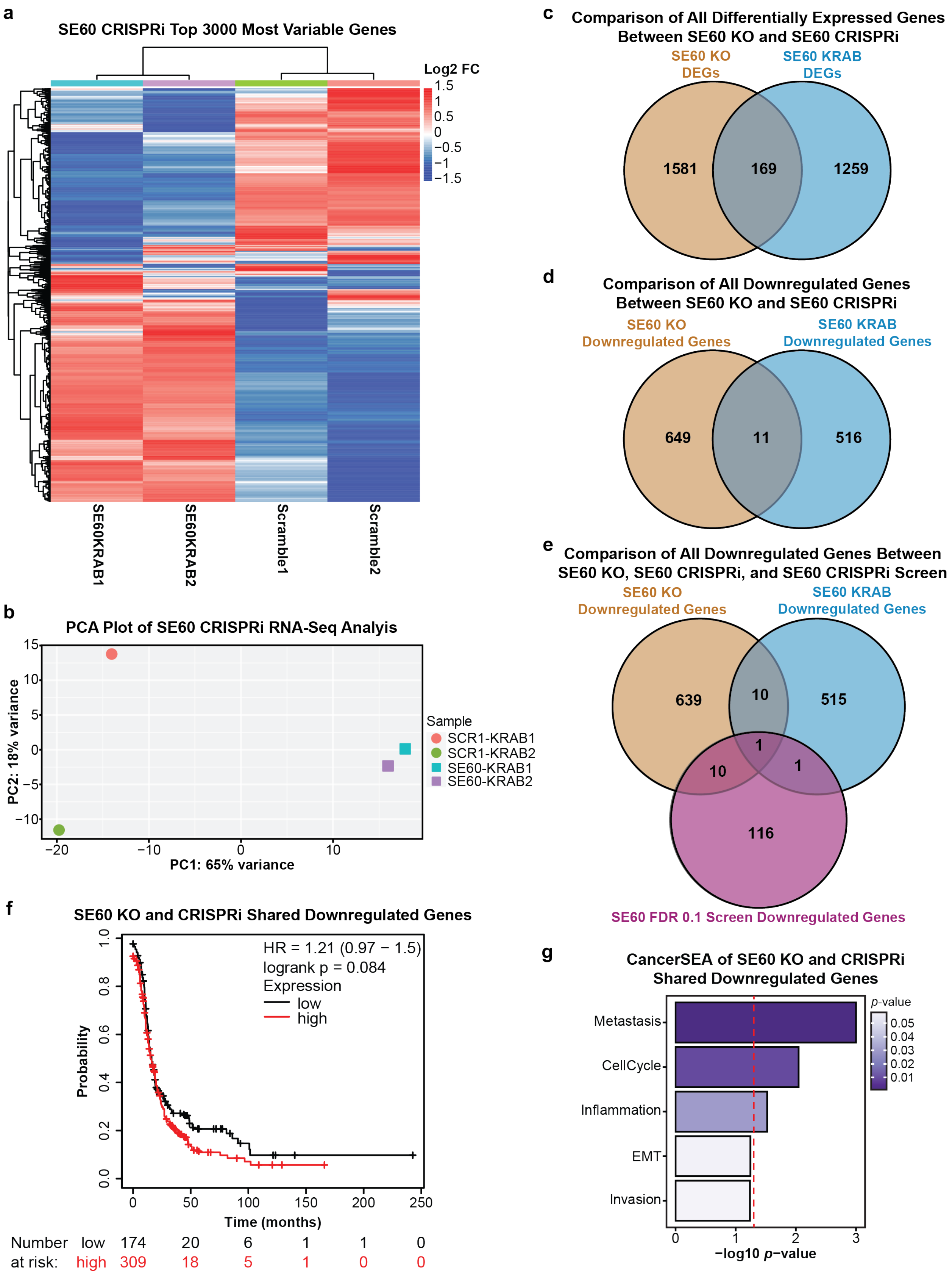

a
SE60 CRISPRi Top 3000 Most Variable Genes
Log2 FC
SE60KRAB1
SE60KRAB2
Scramble1
Scramble2
b
PCA Plot of SE60 CRISPRi RNA-Seq Analyis
PC2: 18% variance
PC1: 65% variance
Sample
SCR1-KRAB1
SCR1-KRAB2
SE60-KRAB1
SE60-KRAB2
c
Comparison of All Differentially Expressed Genes Between SE60 KO and SE60 CRISPRi
SE60 KO DEGs
SE60 KRAB DEGs
1581
169
1259
d
Comparison of All Downregulated Genes Between SE60 KO and SE60 CRISPRi
SE60 KO Downregulated Genes
SE60 KRAB Downregulated Genes
649
11
516
e
Comparison of All Downregulated Genes Between SE60 KO, SE60 CRISPRi, and SE60 CRISPRi Screen
SE60 KO Downregulated Genes
SE60 KRAB Downregulated Genes
639
10
515
1
10
1
116
SE60 FDR 0.1 Screen Downregulated Genes
f
SE60 KO and CRISPRi Shared Downregulated Genes
HR = 1.21 (0.97 − 1.5)
logrank p = 0.084
Expression
low
high
Probability
Time (months)
Number at risk: low 174 20 6 1 1 0
high 309 18 5 1 0 0
g
CancerSEA of SE60 KO and CRISPRi Shared Downregulated Genes
p-value
Metastasis
CellCycle
Inflammation
EMT
Invasion
−log10 p−value

**Supplemental Figure 5: Comparison Between CRISPRi RNA-seq and CRISPR-KO RNA-seq for SE60 Further Suggests Involvement in Metastasis**

- **a.** Unsupervised hierarchical clustering heatmap of the top 3000 most variable genes between scramble controls and SE60 CRISPRi cells measured by RNA-seq.
- **b.** PCA plot showing the variance landscape of both scramble controls and the two SE60 CRISPRi RNA-seq replicates.
- **c.** Overlap of all DEGs detected by CRISPR-KO (left) and CRISPRi (right) targeting SE60; CRISPRi DEGS detected at cutoffs +/- 0.5 Fold Change and adjusted *p*-value of 0.05.
- **d.** Overlap of all downregulated DEGs (potential targets) detected by CRISPR-KO (left) and CRISPRi (right) targeting SE60.
- **e.** Overlap of all downregulated DEGs (potential target genes) detected by CRISPR-KO (left) and CRISPRi (right) targeting SE60 as well as the CRISPRi Screen (bottom).
- **f.** Kaplan Meier plot showing the clinical significance (hazard) of the SE60 KO/CRISPRi shared downregulated DEGS (as noted in D), the red line denotes patients with high expression of this gene signature, significant *p*-values are denoted in red text.
- **g.** CancerSEA gene pathway analysis of the SE60 KO/CRISPRi shared downregulated DEGS; the red line denotes the metric for a *p*-value of 0.05 (significance) converted into the -log10 scale, anything past the line is a significant term.

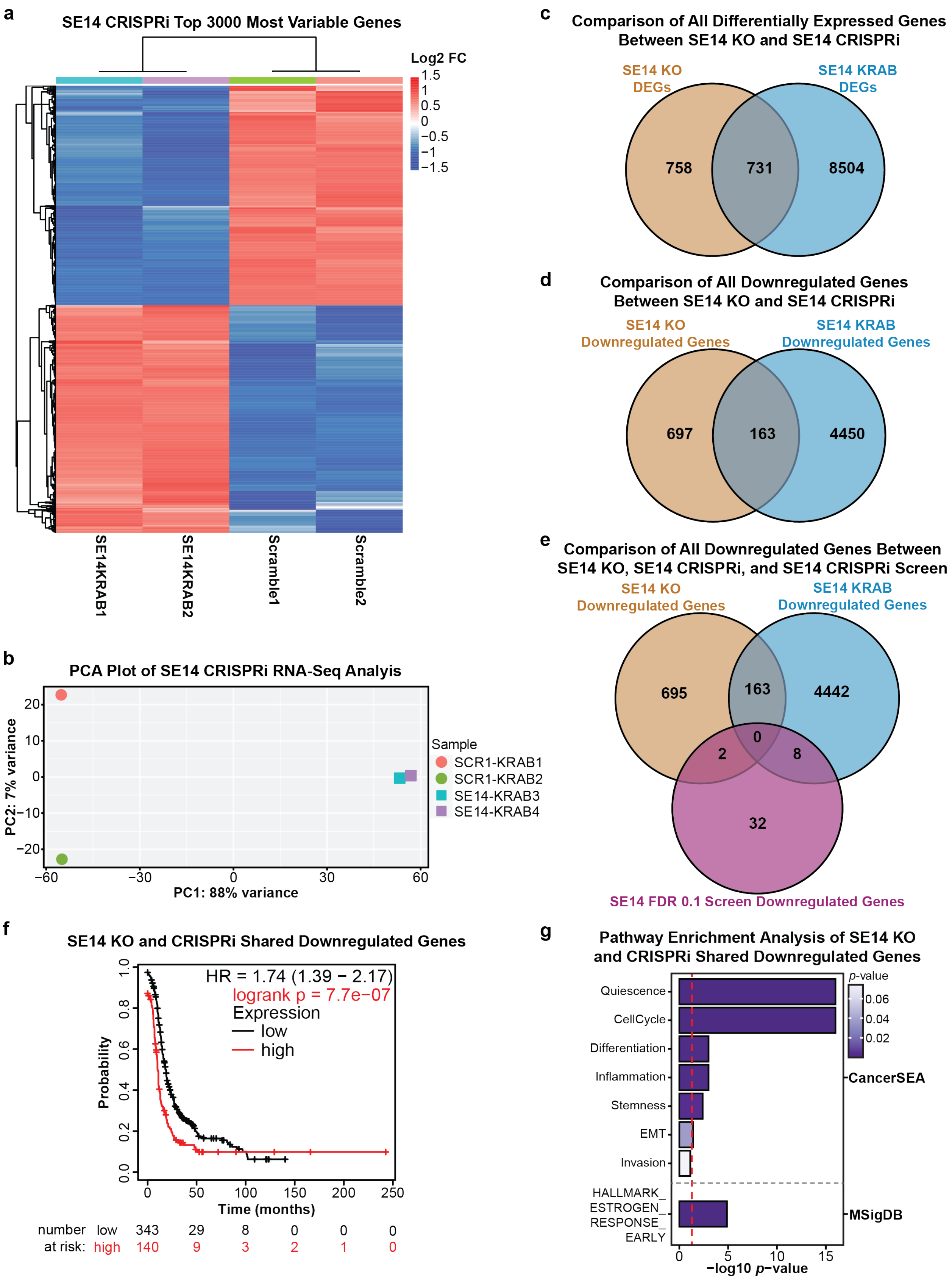
a
SE14 CRISPRi Top 3000 Most Variable Genes
Log2 FC
1.5
1
0.5
0
−0.5
−1
−1.5
SE14KRAB1
SE14KRAB2
Scramble1
Scramble2
b
PCA Plot of SE14 CRISPRi RNA-Seq Analyis
PC2: 7% variance
PC1: 88% variance
Sample
SCR1-KRAB1
SCR1-KRAB2
SE14-KRAB3
SE14-KRAB4
c
Comparison of All Differentially Expressed Genes Between SE14 KO and SE14 CRISPRi
SE14 KO DEGs
SE14 KRAB DEGs
758
731
8504
d
Comparison of All Downregulated Genes Between SE14 KO and SE14 CRISPRi
SE14 KO Downregulated Genes
SE14 KRAB Downregulated Genes
697
163
4450
e
Comparison of All Downregulated Genes Between SE14 KO, SE14 CRISPRi, and SE14 CRISPRi Screen
SE14 KO Downregulated Genes
SE14 KRAB Downregulated Genes
695
163
4442
0
2
8
32
SE14 FDR 0.1 Screen Downregulated Genes
f
SE14 KO and CRISPRi Shared Downregulated Genes
HR = 1.74 (1.39 − 2.17)
logrank p = 7.7e−07
Expression
low
high
Probability
Time (months)
number at risk: low 343 29 8 0 0 0
high 140 9 3 2 1 0
g
Pathway Enrichment Analysis of SE14 KO and CRISPRi Shared Downregulated Genes
Quiescence
CellCycle
Differentiation
Inflammation
Stemness
EMT
Invasion
HALLMARK_ESTROGEN_RESPONSE_EARLY
p-value
0.06
0.04
0.02
CancerSEA
MSigDB
−log10 p−value

**Supplemental Figure 6: Comparison Between CRISPRi RNA-seq and CRISPR-KO RNA-seq for SE14 Further Suggests Involvement in Metastasis**

- **a.** Unsupervised hierarchical clustering heatmap of the top 3000 most variable genes between scramble controls and SE14 CRISPRi cells measured by RNA-seq.
- **b.** PCA plot showing the variance landscape of both scramble controls and the two SE14 CRISPRi RNA-seq replicates
- **c.** Overlap of all DEGs detected by CRISPR-KO (left) and CRISPRi (right) targeting SE14; CRISPRi DEGS detected at cutoffs +/- 0.5 Fold Change and adjusted *p*-value of 0.05.
- **d.** Overlap of all downregulated DEGs (potential targets) detected by CRISPR-KO (left) and CRISPRi (right) targeting SE14.
- **e.** Overlap of all downregulated DEGs (potential target genes) detected by CRISPR-KO (left) and CRISPRi (right) targeting SE14 as well as the CRISPRi Screen (bottom).
- **f.** Kaplan Meier plot showing the clinical significance (hazard) of the SE14 KO/CRISPRi shared downregulated DEGS (as noted in D), the red line denotes patients with high expression of this gene signature, significant *p*-values are denoted in red text.
- **g.** CancerSEA gene pathway analysis of the SE14 KO/CRISPRi shared downregulated DEGS; the red line denotes the metric for a *p*-value of 0.05 (significance) converted into the -log10 scale, anything past the line is a significant term.

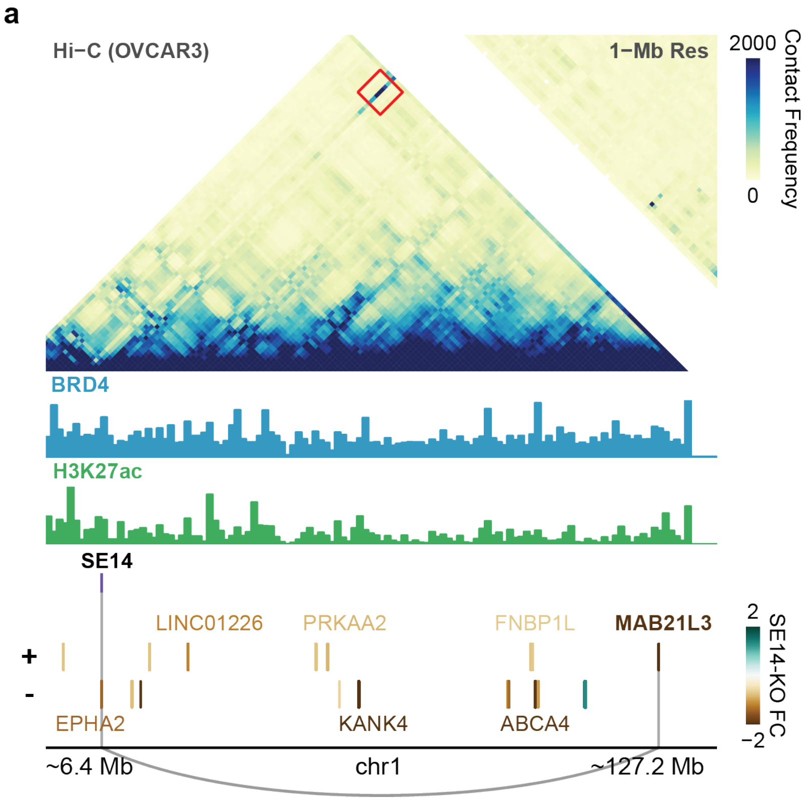


**Supplemental Figure 7: Hi-C Analysis Identifies Strong Long Range Direct Interaction Between SE14 and *MAB21L3***

a. Hi-C contact heatmap showing the interaction between *MAB21L3*, a direct target of SE14, and the SE locus itself (*red square*) across a vast distance.

**a**

**Comparison of SE60-KO Downregulated Genes and Possible SE60 CNVeQTL Genes**

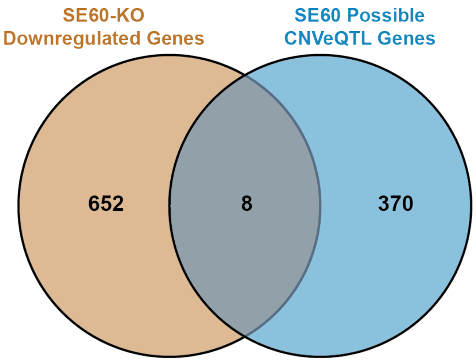


**c**

**Comparison of SE14-KO Downregulated Genes and Possible SE14 CNVeQTL Genes**

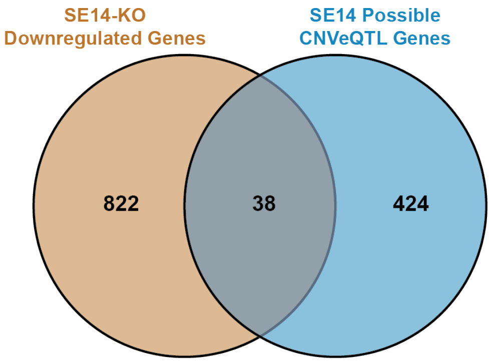


**b**

| SE60 CNVeQTL Genes with Direct Hi-C Contacts (Limited to cis Interactions) |
|---|
| RAE1 |

**d**

| SE14 CNVeQTL Genes with Direct Hi-C Contacts (Limited to cis Interactions) |
|---|
| SKI |
| CLIC4 |
| EPHA2 |
| GNB1 |
| MAP3K6 |
| ACOT11 |
| EDARADD |
| RAP1GAP |

**Supplemental Figure 8: Hi-C Validated *cis* Direct Target Genes are Validated by CNVeQTL Predictions**

- **a.** Venn diagram depicting the results of the overlap analysis between Hi-C validated *cis* direct target genes detected from SE60 KO and all possible *cis* CNVeQTLs.
- **b.** Table showing the results of overlapping the shared cis genes with HI-C cis-direct targets, one gene RAE1 was predicted via CNVeQTL analysis
- **c.** Venn Diagram depicting the results of the overlap between cis-genes detected from the SE14 KO and all possible cis-CNVeQTLs.
- **d.** Table showing the results of overlapping the shared cis genes with HI-C cis-direct targets, eight genes were predicted via CNVeQTL analysis

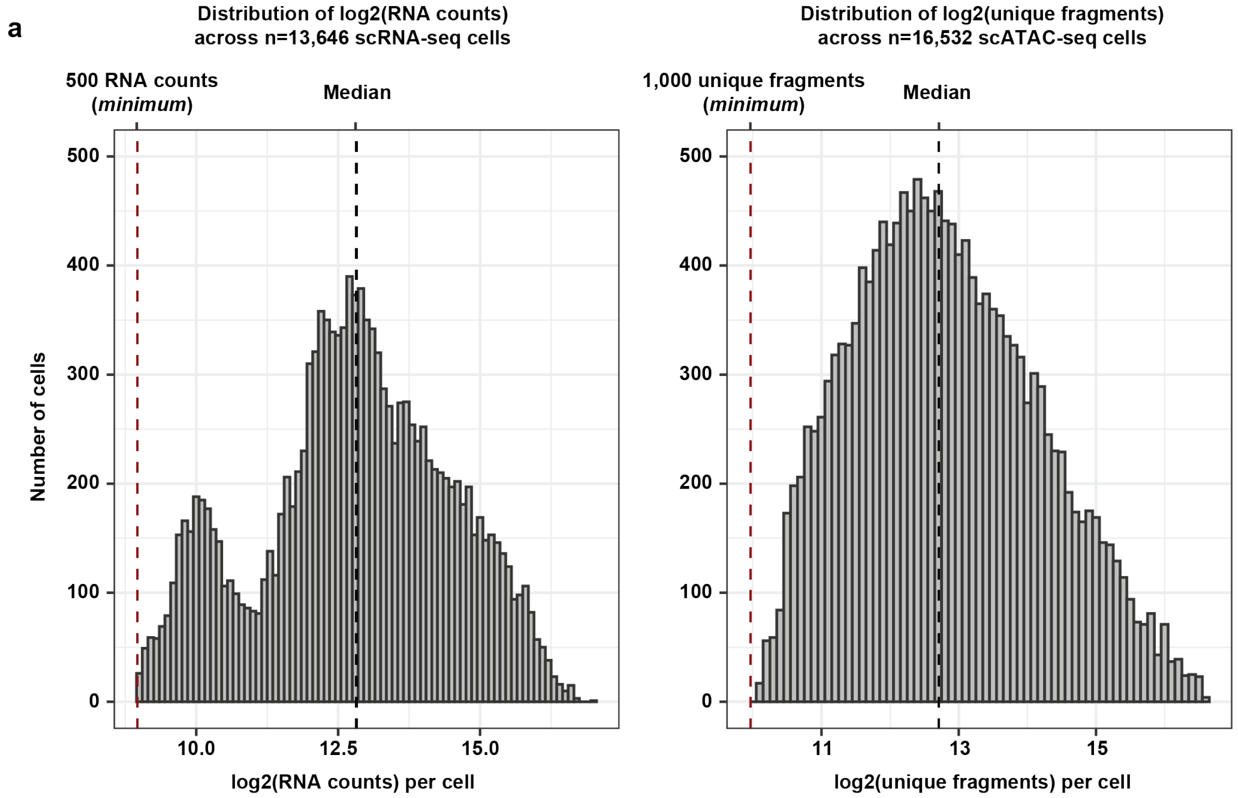


**Supplemental Figure 9: High Quality scRNA-seq and scATAC-seq Data in Terms of RNA Counts Per Cell and Unique Fragments Per Cell**

**a.** Histogram of log2(RNA counts) across 13,646 scRNA-seq cells from both patient samples (*left*). Histogram of log2(unique fragments) across 16,532 scATAC-seq cells from both patient samples (*right*). The dashed red lines represent the standard minimums of 500 RNA counts per cell and 1,000 unique fragments per cell in scRNA-seq and scATAC-seq, respectively. The dashed black lines represent the median values of RNA counts per cell (median=7,234) and unique fragments per cell (median=6,707) in scRNA-seq and scATAC-seq, respectively.

| KO guide target | Forward Sequence | Reverse Sequence | Genomic Coordinates |
|---|---|---|---|
| SE14_LFT_INNR | CACCGAGCGCCTTTCTGAGCGTCAC | AAACGTGACGCTCAGAAAGGCGCTC | chr1:16,499,731-16,499,750 |
| SE14_LFT_OUTR | CACCGACCCTTCAAGTGACCTCGGC | AAACGCCGAGGTCACTTGAAGGGTC | chr1:16,499,554-16,499,573 |
| SE14_RGT_OUTR | CACCGTGCTAAAAGGCTCCCGTAGT | AAACACTACGGGAGCCTTTTAGCAC | chr1:16,502,388-16,502,407 |
| SE14_RGT_INNR | CACCGATTTGAGACCTAATCAGCGG | AAACCCGCTGATTAGGTCTCAAATC | chr1:16,502,255-16,502,274 |
| SE60_LFT_OUT | CACCGTTACACAGAAGGCCGTTCAC | AAACGTGAACGGCCTTCTGTGTAAC | chr20:52,355,052-52,355,071 |
| SE60_LFT_INNR | CACCGAAGTGCAGCTCAACTGCGCA | AAACTGCGCAGTTGAGCTGCACTTC | chr20:52,355,148-52,355,167 |
| SE60_RGT_INNR | CACCGATCGAACTCCTCCCTCGGGA | AAACTCCCGAGGGAGGAGTTCGATC | chr20:52,356,832-52,356,851 |
| SE60_RGT_OUTR | CACCGGTGAAATCATGTATACGAC | AAACGTCGTATACATGATTTCACC | chr20:52,356,874-52,356,893 |

**Supplemental Table 1. Oligonucleotide sgRNA sequences for CRISPR Cas9 Super-Enhancer Deletion.** Sequences are listed 5’ to 3’.

| Primer Name | Sequence |
|---|---|
| SE14-Internal-F | GGTGAAGGGAGAATAGCGGTAG |
| SE14-Internal-R | CTGACTGCCTGATGGGGTTG |
| SE14-External-F | GTGGCTCACCCTTGTAATCTCA |
| SE14-External-R | AGAGAGAGAGAGACAGAGGCAG |
| SE60-Internal-F | GCAGCTCATGATCTCACCAGAG |
| SE60-Internal-R | AGAAGCAAACACTGAAAGCCAC |
| SE60-External-F | TCCCGCATCATTTTCACACATG |
| SE60-External-R | ATTTCCTTCTTCCCAGGCACAA |

**Supplemental Table 2. Super-Enhancer Deletion Screening Primers**. Sequences are listed 5' to 3'.
