## Supplementary material for "A multi-omic dissection of super-enhancer driven oncogenic gene expression programs in ovarian cancer": List_of_Supplemental_Data

### Description of Additional Supplemental Data

**Filename: Supplemental_Data_1.xlsx**
Description: This file contains all of the SE hg19 coordinates, as well as all CRISPRi sgRNA sequences and CRISPRi Screen cluster identity.

**Filename: Supplemental_Data_2.txt**
Description: This file contains all of the hg19 H3K27ac peak coordinates

**Filename: Supplemental_Data_3.txt**
Description: This file contains all of the hg19 BRD4 peak coordinates

**Filename: Supplemental_Data_4.txt**
Description: This file contains all of the co-accessible H3K27ac/BRD4 peaks used to identify SEs via the ROSE pipeline

**Filename: Supplemental_Data_5.txt**
Description: This file contains all of the CNVeQTL identified using Matrix eQTL with a pvalue cutoff of 1e-3

**Filename: Supplemental_Data_6.xlsx**
Description: This file contains all of the genes identified as differentially expressed via the CRISPRi Screen analysis at an FDR of 0.1

**Filename: Supplemental_Data_7.xlsx**
Description: This file contains all of the DEGs detected by DESEQ2 in all CRISPRKO/i experiments

**Filename: Supplemental_Data_8.txt**
Description: This file contains all of the cis-DEGs detected from the CRISPR-KO analysis with information relaying whether or not they are direct or indirect targets as determined by Hi-C
